# Population-Specific Transcriptomic Shifts Underlie Secondary Metabolic Diversification in *Aspergillus flavus* and the Domestication of *Aspergillus oryzae*

**DOI:** 10.1101/2025.10.02.680074

**Authors:** Milton T. Drott, Yen-Wen Wang, E. Anne Hatmaker, Nayanna M. Mercado-Soto, J. Mitch Elmore, Dianiris Luciano-Rosaro, Jae-Hyuk Yu, Antonis Rokas, John G. Gibbons, Anna Huttenlocher, Kunlong Yang, Nancy P. Keller

## Abstract

The genomic signatures of microbial domestication remain poorly understood within the context of natural population variation. Here, we demonstrate that *Aspergillus oryzae*, the filamentous fungus used in soy sauce production, shares more recent ancestry with a predominantly northern, largely non-aflatoxigenic population of *Aspergillus flavus* (population C). Strikingly, *A. oryzae* isolates also overlap with the recently described, clinically enriched *A. flavus* population D, suggesting the possibility of multiple domestication events. Although *A. oryzae* exhibits reduced virulence compared to *A. flavus*, all isolates tested retained pathogenicity in a zebrafish infection model. At the transcriptomic level, *Aspergillus* populations are significantly differentiated, with distinct responses to population density, indicating that population-specific transcriptomes adapt differently to ecological conditions. These differences extend beyond gene content and are not always explained by phylogenetic relationships, suggesting that phenotypic diversification occurs through the rapid reorganization of transcriptomic architectures. For example, *A. oryzae* displays significantly elevated expression of a module enriched for carbohydrate metabolism. Population-specific variation is also evident among secondary metabolite (SM) gene clusters. While *A. oryzae* shows markedly reduced expression of specific SM genes, particularly those involved in aflatoxin biosynthesis, this trend does not extend across the entire secondary metabolome. Using machine-learning-based gene regulatory network inference, we identified population-specific transcriptomic differences linked to distinct transcription factors, with evidence for both cis- and trans-acting regulatory divergence, but no changes in global regulators such as *laeA*. Together, these findings provide new insights into the domestication of *A. oryzae*, its global significance, and the microevolution of fungal secondary metabolic pathways.

**Significance Statement:** *Aspergillus flavus* poses a significant threat to global agriculture by infecting crops and producing aflatoxin, a potent carcinogen that undermines food safety and international trade. The species is also a major human pathogen causing invasive fungal infections. Here, we demonstrate that *A. flavus* populations exhibit not only variations in their capacity to produce toxins but also in the way they regulate gene expression in response to environmental cues. These regulatory differences played a key role in the domestication of *Aspergillus oryzae*, a non-aflatoxigenic relative that has been safely used in food fermentation for centuries. Notably, while *A. oryzae* exhibits reduced aflatoxin biosynthetic gene expression, it expresses other secondary metabolite genes and retains limited opportunistic pathogenic potential. By revealing how transcriptomic rewiring drives population-level divergence, this work provides new insights into the evolution of toxin production, with broad implications for microbial domestication, crop protection, and global food security.

## Introduction

The domestication of plants and animals is often cited as a key factor in explaining the success of burgeoning human civilizations. The impact of domestication on specific animal traits has been studied since Darwin (Darwin, 1868), leading to the definition of ‘domestication syndromes,’ that define similar patterns of co-occurring traits across domesticated organisms. Similar patterns have also been described among domesticated plants (e.g., barley, diploid einkorn wheat, and tetraploid emmer wheat (Brown et al., 2009)). While the phenotypes of domestication have been well described, disentangling the evolutionary histories of these organisms has often required insights from genomics. Recent pangenomic studies have demonstrated that dogs in the Near East and in Africa are descended from a specific population of Eurasian wolves, suggesting admixture or an independent domestication event (Bergström et al., 2022). Clarifying the ancestry of domesticated populations is essential for identifying how differences in gene content or genetic differentiation may contribute to phenotypic differentiation (Gong et al., 2023). Despite progress in understanding the evolution of charismatic or economically important domesticated plants and animals, microbial domestication has been relatively overlooked (Steensels et al., 2019). Domesticated microbes, like plants and animals, have shaped human history and the way we eat today. These organisms, however, do not always exhibit the fundamental characteristics of domestication syndromes that are evident in most plant and animal species. Indeed, Rokas (Rokas, 2009), speculated that a lack of relaxed selection acting on some domesticated microbes may reflect fundamental differences in population dynamics of microbes.

*Aspergillus oryzae* is thought to have been domesticated thousands of years ago: the solid-state cultivation techniques used to culture this fungus date back 2,000- 3,000 years ago (Murakami, 1980) and the first commercial sales of Koji (rice fermented with *A. oryzae*) are evident in the 13-15^th^ century (Machida et al., 2008). Early descriptions of *A. oryzae* listed it as a variety of *A. flavus* (Wehmer, 1901), but by 1921, it was recognized as a separate species (Thom & Church, 1921). Molecular evidence appears unequivocal that *A. oryzae* was domesticated from *Aspergillus flavus*, a plant and animal pathogen notorious for its production of the potent carcinogen, aflatoxin. Gibbons et al (Gibbons et al., 2012) clarified that *A. flavus* isolates span a spectrum of relatedness to *A. oryzae* and suggested that domestication likely occurred from an atoxigenic lineage of *A. flavus*. Balancing selection for aflatoxin production (Carbone et al., 2007; Drott et al., 2017; Drott, Debenport, et al., 2019) is thought to maintain closely related aflatoxigenic and non-aflatoxigenic lineages. The fact that aflatoxigenic and non-aflatoxigenic isolates are often found at similar prevalences in the same geographic area (Drott et al., 2020) raises a question of how this organism could have been domesticated without many poisonings (although this human cost could have been lost to history). Intriguingly, the population density of *A. flavus* decreases dramatically at higher latitudes (Drott, Fessler, et al., 2019), and there’s evidence of a largely northern population that, in contrast to other populations, is almost entirely non-aflatoxigenic (Drott et al., 2020). Notably Japan, where Koji was first produced (Machida et al., 2008) commercially, is predominantly northern. To clarify the impact of domestication on *A. oryzae*, there is a need to examine extant relatives that more closely reflect the lineage originally used for domestication.

Microbes, just like plants and animals, are domesticated through artificial selection of desired traits. In contrast to macroscopic organisms, selection on microbes is typically imposed for their chemical diversity. Gibbons et al. (Gibbons et al., 2012) found selection in *A. oryzae* acting on flavor-related glutaminase genes and revealed an increased copy number of amylase genes. Critically, artificial selection would also strongly select against toxic metabolites. A hallmark of the domestication of *A. oryzae* has been a dramatic shift in the expression of several secondary metabolite (SM) gene clusters that often encode toxic compounds (including aflatoxin) were downregulated (Gibbons et al., 2012). Transcriptomic rewiring is thought to be a major mechanism of phenotypic differentiation among even very closely related species, e.g. humans and other apes (Gould, 1977; Johnson, 2017). The expression of *Aspergillus* SMs is impacted by ecological factors that domestication would have shifted, including nutrient environment (Georgianna et al., 2010) and population density (Affeldt et al., 2012). Density-dependent transcriptomic rewiring is known to suppress aflatoxin production at higher densities (Affeldt et al., 2012). Nearly 25% of the pan-secondary metabolome in *A. flavus* is differentiated between populations (Drott et al., 2021). However, almost all past transcriptomic work in *A. flavus* has focused on a single reference genome, leaving questions about the population-transcriptomic patterns that may clarify the domestication of *A. oryzae*.

To investigate the molecular basis of population-level variation, we conducted a comparative transcriptomic analysis of *A. flavus*. Specifically, we sought to determine whether: (i) transcriptomic profiles are structured by population-specific differentiation; (ii) distinct populations exhibit divergent responses to density-dependent metabolic rewiring; (iii) forces beyond phylogenetic relatedness have shaped the evolution of transcriptomic architectures; and (iv) population-specific transcriptional programs can be attributed to a discrete set of regulatory factors. We also investigate the impact of domestication on the virulence of *A. oryzae*. Together, these analyses reveal the mechanisms by which regulatory divergence contributes to metabolic diversification in *A. flavus*, providing new insights into the domestication trajectory of *A. oryzae*.

## Methods

### Population sampling and genome sequencing

We sampled five to seven isolates from the three *A. flavus* populations that were known at the inception of this project (Drott et al., 2020). From each population, a mixture of aflatoxigenic and non-aflatoxigenic isolates were selected as evenly as possible (Table S1) while sampling broadly from across each population’s clade (based on results from Drott et al. (2021)). An additional six *A. oryzae* isolates (Table S1) were collected from public and author’s culture collections. *A. oryzae* isolates were similarly chosen to be as genetically and geographically as diverse as possible, although little information was typically available for isolates and industrial strains are rarely shared. Of these, two (RIB40 and NRRL506) already had genome sequences available. We sequenced the genomes of the remaining four isolates using methods similar to Drott et al. (2020) using the Illumina NovoSeq 6000 platform. Resulting sequence data was used to call SNPs using the GATK (Van der Auwera & O’Connor, 2020) pipeline as defined by Drott et al. (2020). Resulting SNPs were integrated with data from Drott et al. (2020) and used to construct a neighbor-net network in SplitsTree v6.3.31 (Huson & Bryant, 2006) to confirm that isolates used in the study were not clones (see methods below). While NRRL506 shared recent common ancestry with RIB40, it was retained given the distant isolation locations of the two isolates and questions about the importance of clonality in this domesticated lineage (Figure S1). A final list of selected isolates is available in Supplemental Table S1.

### Acquisition of public genomes, annotation, and reference selection

All genomes of *A. flavus* and *A. oryzae* that were available on NCBI were downloaded on 5/24/22 (a list of accessions used here are available in Table S2). Genomes were annotated with AUGUSTUS v3.3.2. with pre-built *A. oryzae* gene models (Stanke & Morgenstern, 2005). A chromosome assembly of the *A. flavus* reference genome, NRRL3357 (Aspfl2_3), and the associated genome-annotation that was originally developed in Drott et al. (2020) was obtained from JGI was used as a reference genome.

### Pangenome construction

We organized predicted proteins from our total genomic dataset into orthogroups using OrthoFinder v2.5.5 (Emms & Kelly, 2019). Functional annotations of proteins in orthogroups were identified from PFAM domain-associated GO terms using InterProScan v5.44-79 (Jones et al., 2014). Orthogroups found in all genomes (a stringent cutoff) were considered core.

### Analysis of SM gene clusters

Biosynthetic gene clusters (BGCs) were identified from genome assemblies using antiSMASH v5.1.2 (Blin et al., 2019). We obtained manually curated borders of 15 characterized SM BGCs and the sequences of population-specific BGCs from Drott et al. (2021). The relationship of BGCs showing population-specific variations were identified compared to Drott et al. (2021) using BLASTp v2.12.0+ from the BLAST+ suite (Camacho et al., 2009). BGCs were dereplicated by creating networks with Biosynthetic Gene Similarity Clustering and Prospecting Engine (BiG-SCAPE) (Navarro-Muñoz et al., 2020), which were visualized with Cytoscape v3.7.1 (Su et al., 2014). BGCs that did not map to the reference genome or to other BGCs defined in Drott et al. (2021) are called “combos”. A table of clusters mapped to the reference genome is available in Table S3.

### Phylogenetic tree reconstruction

The original sampling of isolates was based on population structuring inferred by Drott et al. (2021). To confirm these relationships in the presence of public data, we reconstructed phylogenetic trees from assembled genome sequences. Benchmarking single copy ortholog (BUSCO) sequences corresponding to the fungal database v10 were obtained using BUSCO (Simão et al., 2015). BUSCO-based Maximum likelihood phylogenies were constructed by aligning sequences with MAFFT (Katoh et al., 2002), trimming with trimAL (Capella-Gutiérrez et al., 2009), and using the test parameter across a partition model in IQ-TREE2 (Nguyen et al., 2015) with pre-filtering similar to previous work (Steenwyk et al., 2019) as is detailed in the Supplemental Methods. Phylogenetic trees were also constructed for each of the genes in the Eurotiales BUSCO dataset and used to construct a quartet-based tree in ASTRAL (Zhang et al., 2018). We also constructed phylogenies from SNP data using IQ-TREE2 with methods identical to those used previously (Drott et al., 2021). Additionally, SNP data from Hatmaker et al. (2025) was integrated with data from this study using IQ-TREE2, as detailed in the supplemental methods. Resulting trees were visualized using ggtree (Yu et al., 2017) and iTOL v6 (Letunic & Bork, 2024), respectively.

### Clone Correction

To collapse genetically identical individuals, we performed clone correction similar to DeGenring et al. (2025). Briefly, the ‘mldist’ data matrix output by IQ-TREE2 from the BUSCO maximum likelihood tree was reformatted into a list of pairwise comparisons between isolates using a custom script that uses the R package *reshape2* v1.4.4 (Wickham, 2007). The resulting pairwise distances were visualized as a histogram. We assumed that the peak and tail at/near zero genetic distance represents true clonal relationships, and a distance cutoff was selected to capture this near-zero distribution. When isolates shared a pairwise distances less than our cutoff, they were collapsed into a single multi-locus genotype (MLG, Table S2, Figure S2). When clone corrected data were used, the genome with the highest BUSCO completeness metric was selected to represent an MLG. This approach is flexible across datasets, allowing for data-driven and transparent clone correction of diverse datasets.

### Experimental design and population transcriptomic sequencing

To dissect the transcriptomic differentiation of *A. flavus* and *A. oryzae* populations, we chose an experimental design that favored biological replication within population instead of technical reps, using 5-7 isolates from each population as biological replicates within population (Table S1) with two technical replicates each.

Fungal spores were harvested from colonies growing on PDA using 0.05% tween 20 water. Resulting spore suspensions were diluted to either 10^2^ or 10^6^ spores/ml and a 200 ul suspension was then spread evenly across a piece of sterile cellophane resting on a PDA. After 100 h, mycelium was collected, frozen in liquid nitrogen, and lyophilized for 24 h. Two technical replicates for each plate were harvested and combined and extracted using trizol (Ambion, USA), according to the manufacturers protocol. RNA purity and integrity were tested via gel electrophoresis, nanodrop and in the Agilent 2100 bioanalyzer. RNA concentration was quantified precisely by Qubit 2.0. Libraries were prepared using the poly-A mRNA enrichment protocol and paired-end sequencing in the NovaSeq 6000 platform.

### Identification of differentially expressed genes

The transcriptomic read count of each gene was quantified with kallisto ver. 0.44.0 (Bray et al., 2016). A Wald test from the R package DESeq2 (Love et al., 2014) was used to identify the differentially expressed genes between spore density for each population.

### Co-expression network

Weighted Gene Co-expression Network Analyses (WGCNA) were used to identify co-expressing gene modules at a population level. First, the read counts were normalized with Trimmed Mean of the M-values (TMM) and Fragments Per Kilobase per Million mapped fragments (FPKM) using R package *edgeR* (Robinson et al., 2010). Next, a weighted gene co-expression network was constructed using R package *WGCNA* with the log values of TMM-FPKM (Langfelder & Horvath, 2008). The dendrogram cut height to merge modules was set to 0.25 and the soft threshold was set to 12. To understand the putative functions of these gene modules, R package topGO was used to identify the gene ontology (GO) terms enriched in each module (Alexa & Rahnenfuhrer, 2010).

### Phylogenetic analysis of transcriptomics

To test if genes were differentially expressed in different populations, we used three strategies: (1) PERMANOVA, (2) phylogenetic ANOVA and (3) two-way ANOVA. The function *adonis2* from R package *vegan* (Dixon, 2003) was used to perform a PERMANOVA testing if genome-wide gene expression level (log-TMM-FPKM) is dependent on population, spore density, toxicity, or any interacting terms. The function *aov.phylo* from R package *geiger* (Pennell et al., 2014) was used to perform phylogenetic ANOVA focusing on a single spore density at a time to separate the effects of the independent variables between the two treatments. Two-way ANOVAs were done with base R using population, spore density and their interacting term as independent variables and gene expression level as the dependent variable. The P-values from ANOVA were further adjusted again using False Discovery Rate (FDR) for the number of genes tested. The analyses were also done on gene expression modules (normalized expression value) and the backbone genes of secondary metabolism clusters (log-TMM-FPKM; mean of log-TMM-FPKM if multiple backbone genes were identified), but the toxicity term and any associated interaction terms were not included in the PERMANOVA.

### Gene regulatory network (GRN) inference and analyses

Normalized read counts from the co-expression network analysis were used as inputs to SC-ION v3.2 (https://github.com/nmclark2/SCION) (Clark et al., 2021), which employs a modified version of the tree-based regression method GENIE3 (Huynh-Thu et al., 2010). Networks were inferred separately using only differentially expressed genes from each of the condition, population, and condition*population contrasts (adj. P < 0.05) based on two-way ANOVA analysis (Table S6). Transcription factors annotated in MycoCosm (https://mycocosm.jgi.doe.gov/Aspfl2_3/Aspfl2_3.home.html) were assigned as potential regulators, and all differentially expressed genes were assigned as potential targets. The resulting GRNs were trimmed to remove edge weights < 0.65. This cutoff corresponded to the best supported 0.23%, 1.3%, and 15.7% of edges in the population, density, and population * density contrast networks, respectively. GRNs were analyzed using the *igraph* and *influential* R packages (Csardi & Nepusz, 2006), and node centrality was calculated using the Integrated Value of Influence (IVI) metric (Salavaty et al., 2020). Networks were visualized in Cytoscape version 3.10.2 (Shannon et al., 2003).

### Ethics Statement

The Institutional Animal Care and Use Committee (IACUC) at the University of Wisconsin-Madison College of Agricultural and Life Sciences (CALS) approved the use of zebrafish in this research. The animal care and use protocol M005405-A02 adheres to the guidelines established by the federal Health Research Extension Act and the Public Health Service Policy on the Humane Care and Use of Laboratory Animals, led by the National Institutes of Health (NIH) Office of Laboratory Animal Welfare (OLAW).

### Zebrafish Husbandry and Maintenance

Wild-type zebrafish (*Danio rerio*) lines were maintained under standard conditions as described previously (Knox et al., 2014). For experiments, embryos were collected and maintained at 28.5 °C in E3 water.

### Fungal Inoculum Preparation and Microinjections

Fungal strains were grown from glycerol stocks on Glucose Minimal Media (GMM), and spores were harvested and prepared as previously described (Schoen et al., 2021, 2023). The conidial concentration was adjusted to 1.5 × 10^8^ conidia/mL and then mixed 2:1 with 1% Phenol red before injection. Larvae were anesthetized at 2 days post-fertilization and injected with 3 nL of conidial suspension into the hindbrain ventricle. After injections, larvae were rinsed three times with E3 and transferred to individual wells in 96-well plates. Survival was monitored for 7 days. After injections, eight larvae per condition were transferred to individual microcentrifuge tubes containing 90 μL 1X PBS with 500 μg/mL kanamycin and 500 μg/mL gentamycin. Larvae were homogenized for 15 seconds in a mini bead beater, plated on GMM, and incubated for 2 days at 37 °C. After incubation, colony-forming units (CFUs) were counted and averaged for each condition. The pairwise log-rank test comparisons in larval survival were calculated using the R packages survival v3.8-3 (Therneau, 2015) and survminer v0.5.0 (Kassambara et al., 2016), using the pairwise_survdiff() command, with Benjamini-Hochberg adjustment for false-discovery rate correction.

## Results

### *A. oryzae* Shares Recent Common Ancestry with *A. flavus* Population C and Overlaps with a Clinically Enriched Lineage

To clarify the domestication origins of *A. oryzae*, we reconstructed phylogenetic relationships inferred phylogenies from genome-wide SNP datasets including all 793,820 biallelic SNPs, 629,271 SNPs with no missing data, 430,178 intergenic SNPs, and a thinned (to 1 kb distance) subset of 20,067 intergenic SNPs (Figure S3). We also constructed a neighbor-net network to better infer potential recombination as reticulation (Figure S1). All phylogenetic methods consistently offered strong support that *A. oryzae* shares the most recent common ancestry with population C of *A. flavus*, a largely northern and non-aflatoxigenic lineage. BUSCO-based inferences used to incorporate public gene presence/absence data (discussed below in this section) were sometimes unstable and/or gave different topologies (Figure S4 and S5). SNP datasets are far more robust in resolving intraspecific variation than highly conserved gene sets like BUSCO; we rely on SNP-based relationships for our inferences of population relationships. Overall, we conclude that *A. oryzae* was domesticated from *A. flavus* and shares more-recent common ancestry with population C.

While preparing this manuscript, we became aware of a newly identified *A. flavus* population (population D) enriched for clinical isolates (Hatmaker et al., 2025). We integrated SNP data from our *A. oryzae* isolates into the Hatmaker dataset and found that *A. oryzae* isolates were interspersed within population D (Figure 1A). Nearly all *A. oryzae* isolates fall into a polyphyletic structure of *A. oryzae* around population D, suggesting multiple domestication events and/or recombination. This reanalysis also found that population D and *A. oryzae* were strongly supported as most closely related to population C, the same topology as we identified above (Figure S6). The genomes of population D isolates are significantly larger than those in all other populations, perhaps reflecting a recent expansion (Figure S7). As these data became available only at a late stage of the study, population D was not included in all subsequent analyses, although the results presented here provide important context.

**Figure 1.**
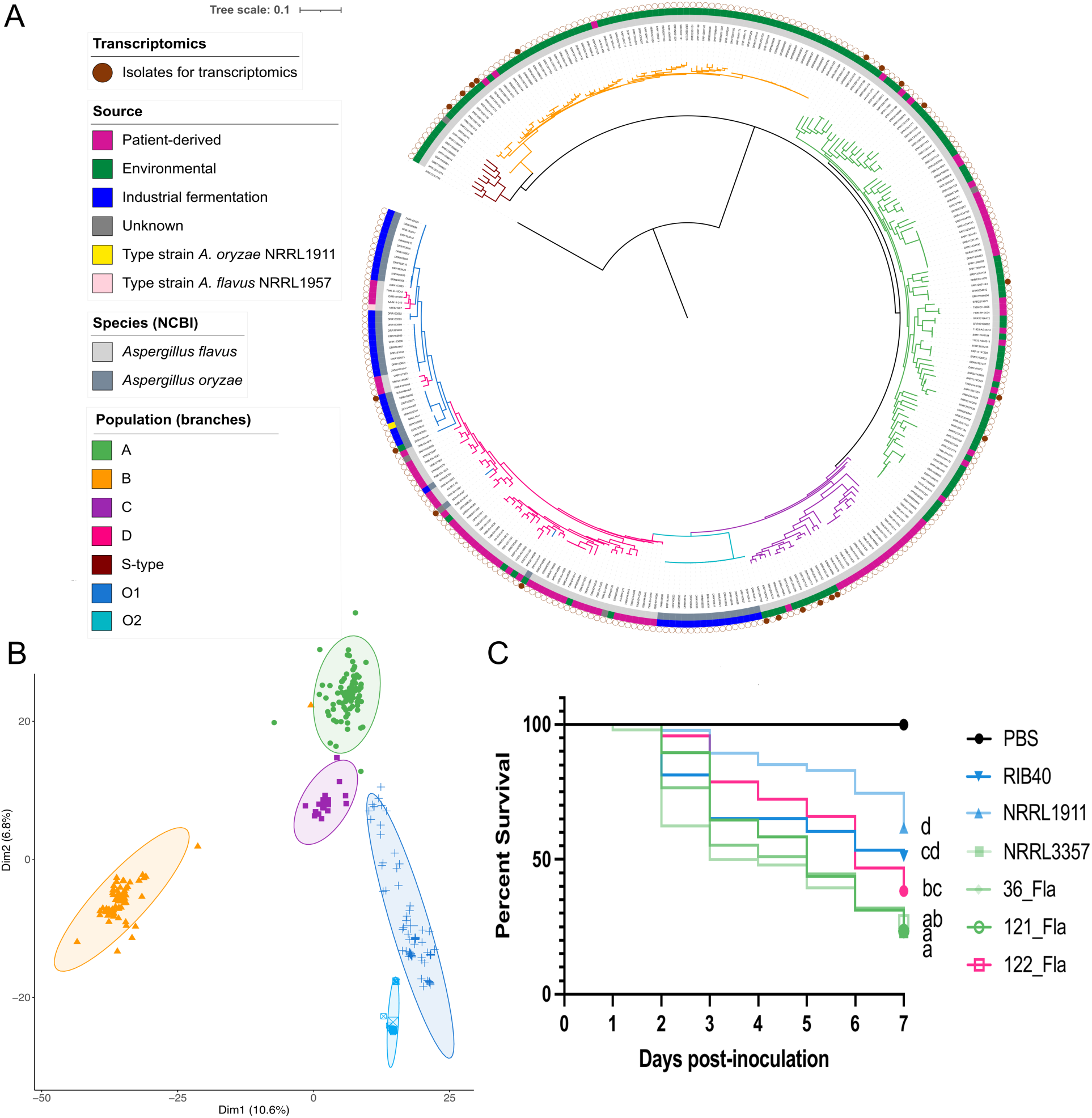
*Aspergillus oryzae* shares recent common ancestry with *Aspergillus flavus* population D and Population C. A) A maximum likelihood phylogeny constructed using genome-wide SNPs including data from Hatmaker et al. (Hatmaker et al., 2025) is rooted at *Aspergillus minisclerotigenes*. Bootstrap supports for this tree are depicted in Figure S6. B) principal component analysis of *A. flavus* isolates across orthogroups showing the relationship in isolates and populations based on gene-space. Colors correspond to populations defined in A. C) Kill curve of zebrafish assay demonstrating that *A. flavus* isolates from population A, D, and *A. oryzae* all show virulence in animal models. Kaplan-Meier survival curve of zebrafish assay demonstrating that *A. flavus* isolates from population A and D, and *A. oryzae* all show virulence in an animal model compared to the PBS control. Lines are colored in shades that correspond to populations defined in panel A. Letters indicate statistical significance groups (log-rank test, P < 0.05, Benjamini-Hochberg adjusted P-values). Average spore dose injected: RIB40 = 26, NRRL1911 = 48, NRRL3357 = 34, 36_Fla = 42, 121 Fla = 46, 122 Fla = 38; n = 43-48 larvae per condition.

Principal component analysis of gene presence/absence data supported the phylogenetic results, with *A. oryzae* grouping most closely with population C (Figure 1B). Functional enrichment among differentially present genes has recently been addressed by Hatmaker et al. (Hatmaker et al., 2025); here we find enrichments includes categories related to reproduction, viruses, and transposable elements (Supplemental Results).

Clonality varied widely across populations: A (27.5%), B (43.7%), and C (26.7%). In contrast, *A. oryzae* showed a higher overall clonal fraction (75.2%) and split into two major clades — designated O1 and O2 — with clonal fractions of 66.7% and 97%, respectively (Figure S2, Table S2). Both clades contained industrial strains, but only O1 included environmental isolates. The O2 clade was composed of just two genetically distinct multi-locus genotypes (Figure S2 and S4).

### *A. oryzae* retains virulence, but less so than *A. flavus* in zebrafish model

The genetic similarities between clinical *A. flavus* isolates and *A. oryzae* led us to explore virulence of these groups using a zebrafish model of aspergillosis (Fig. 1C). We used two *A. oryzae* reference isolates (RIB40 and NRRL1911 [the type strain of the species]), the *A. flavus* reference strain NRRL3357 (population A), and three *A. flavus* clinical isolates (36_Fla, 121_Fla, [both population A] and 122_Fla [population D]). All isolates were able to cause mortality. *A. oryzae* isolates were less virulent than all *A. flavus* isolates (log-rank test, p < 0.0001). The isolates from population A (121_Fla and 122_Fla) were slightly more virulent than the isolate from population D (log-rank test, p = 0.047) and more virulent than either RIB40 (log-rank test, p = 0.028) or NRRL1911 (log-rank test, p < 0.0001). The population D isolate was also significantly more virulent than NRRL1911 (log-rank test, p = 0.028) but not RIB40.

### Population transcriptomics reveal population-specific expression profiles

We examined whether gene expression varied significantly between populations and how this variation was influenced by an ecologically relevant factor: culture density, which influences SM production and fungal development (Horowitz Brown et al., 2008). Gene expression and gene module expression were both significantly differentiated among populations (PERMANOVA, P = 0.000999, Table S4 and S5 respectively), spore density (P = 0.002997;) and their interaction was significant across genes (P = 0.030969), and weakly across modules (P = 0.066933). Aflatoxin producing ability did not have a significant impact on expression profiles. A total of 4,568 genes were differentially expressed among populations (two-way ANOVA, adjusted P < 0.05, Table S6). As expected, culture density also influenced transcriptomic profiles, but its effect was modest, with only 229 genes showing differential expression. Notably, the transcriptomic response to density of 73 genes varied significantly across populations (P < 0.05), suggesting population-specific plasticity. At low density, 21 modules were differentially expressed between populations (Table S7); 10 modules were differentially expressed at high density (ANOVA, unadjusted P < 0.05, Figure 2, Table S8). While population-specific gene content may contribute to these findings, over 70% of differentially expressed genes (DEGs)—across population, condition, and their interaction—resided in the core genome offering robust validation of this patterning as similar studies have not performed tandem pangenomics and transcriptomics.

**Figure 2.**
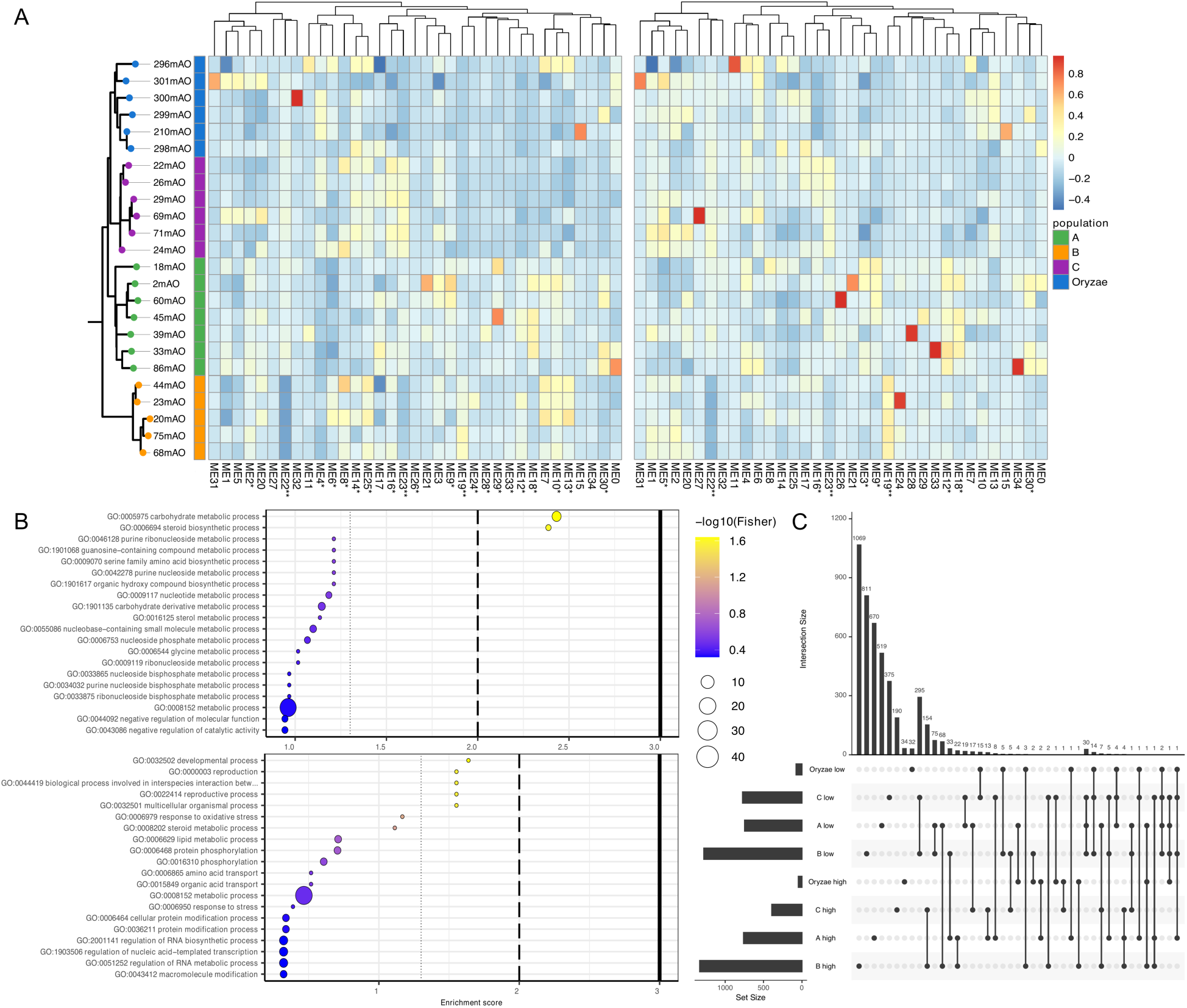
*Aspergillus flavus* and *Aspergillus oryzae* populations differentially express genes and modules as a function of colony density. (A) Heatmaps depicting the expression of module eigenvalues obtained from Weighted Gene Co-expression Network Analyses across isolates grown at low (left) or high (right) density. One star on the module labels represents significant differentiation between populations (ANOVA) while two stars indicates significance when accounting for phylogenetic distances (phylogenetic ANOVA). Isolates of *A. flavus* and *A. oryzae* populations are organized according to a maximum-likelihood phylogeny constructed from the genetic relationships of 173 high-quality BUSCO genes. Hierarchical clustering based on combined density data is presented above both heatmaps. (B) Two examples of the GO enrichment biological process terms present in significantly differentially expressed modules are given for module ME14 (top) and ME22 (bottom). The three increasingly solid dotted lines on enrichment plots indicate significance at the 0.05, 0.01, and 0.001 levels respectively. (C) The differential expression of individual genes is represented in an upset plot that depicts which differentially expressed genes (DEGs) are more highly expressed in a given spore density (e.g., there are 295 DEGs shared between pops B and C that are expressed more in the low spore density and 154 genes that are shared between these pops at the high spore density, for a total of 449 density-dependent DEGs shared between these populations).

Three co-expression modules (19, 22, and 23) showed significantly greater expression divergence than expected from phylogeny alone (phylogenetic ANOVA; adjusted P < 0.05, Table S8). Module 22, which had notably low expression in population B, was enriched for reproduction and development genes (Fisher’s exact test, unadjusted P = 0.023). Module 14 was more highly expressed across all groups compared to population A at low density (ANOVA, P = 0.0226; phylogenetic ANOVA, P = 0.735). At high density, expression was highest in three *A. oryzae* isolates, though population-level differences were nonsignificant (ANOVA, P = 0.149). The latter module was enriched for carbohydrate metabolism genes (Fisher’s exact test, unadjusted P = 0.0037; Figure 2B).

### Population and phylogenetic patterns of SM expression

Previous work suggests that *A. oryzae* does not express certain highly toxic SM BGCs, including aflatoxin. To investigate the evolution of this phenomenon, we examined expression patterns of SM backbone genes across *A. oryzae* and three *A. flavus* populations. Overall SM BGC backbone expression differed significantly between populations, densities, and weakly as an interaction between the two (PERMANOVA, P = 0. 000999, P = 0.006993, and P = 0.057942 respectively, Table S9). While differences in transcriptomic space across the entire genome were poorly defined in a principle component analysis of all genes, analysis of SM backbone genes identified a shift in the transcriptomic space of *A. oryzae* (Figure S8). Hierarchical clustering revealed distinct expression clades: one clade (∼20 backbones, leftmost in Figure 3) was highly expressed across isolates and conditions; another (∼15 backbones, central clade) showed intermediate expression. The rightmost clade had generally low expression with three sub clades (from left to right): one included three backbones—*aflavarin*, *aflatrem* (ATM1), and a *piperazine* backbone—with generally low expression, except in a small subset of isolates from populations B and C; the second subclade universally showed close to no expression and the final subclade showed low expression (Figure 3). Given the dogma that BGCs are silent under laboratory conditions, it is surprising how many backbone genes showed detectible level of expression although we acknowledge that transcript abundance is not necessarily equivalent to metabolite production.

**Figure 3.**
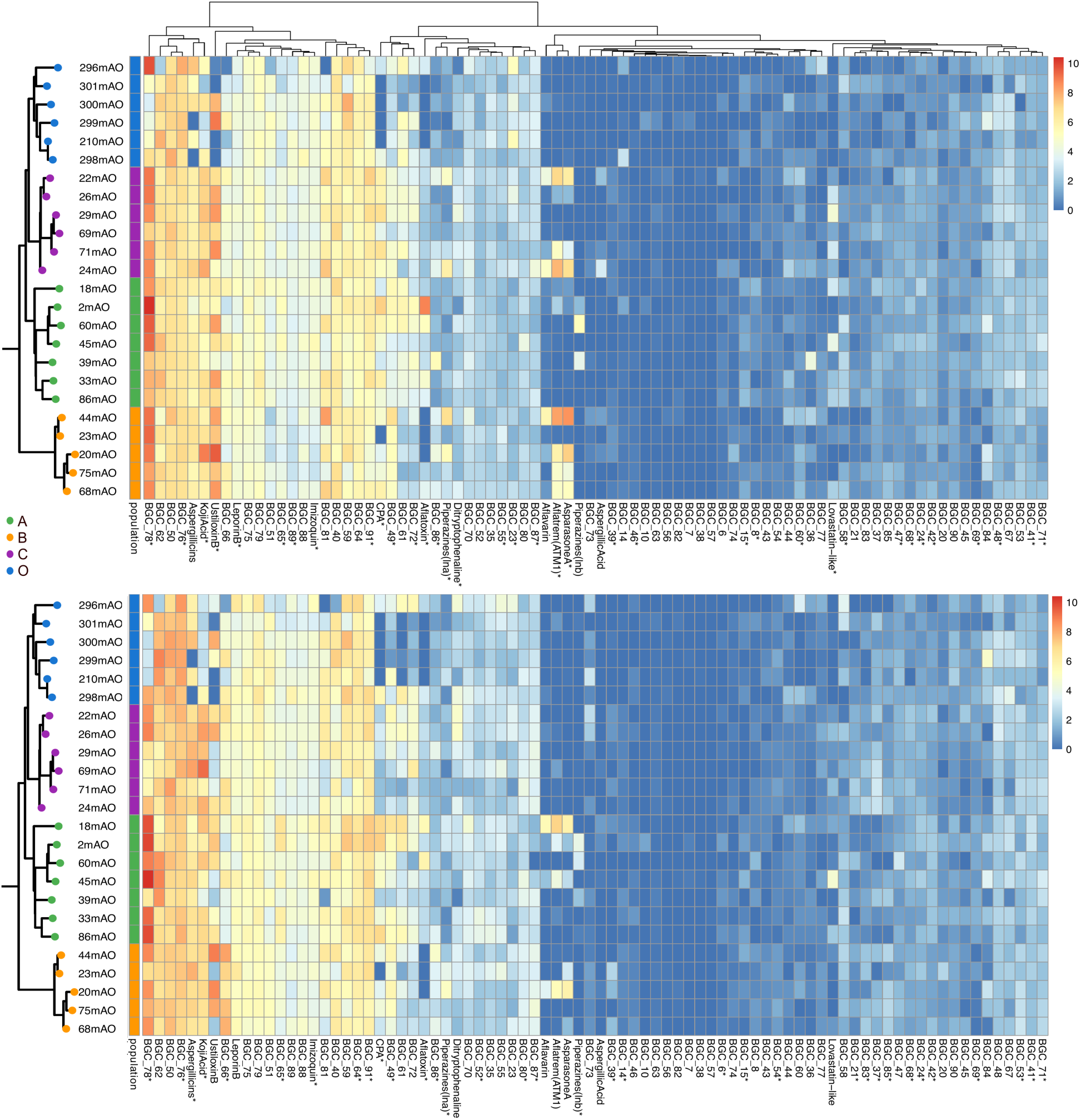
Heatmap depicting the normalized expression counts of secondary metabolite backbone genes that are present in the NRRL_3357 *Aspergillus flavus* reference genome as expressed at low (top) and high (bottom) population density. Isolates of *A. flavus* and *Aspergillus oryzae* populations are organized according to a maximum- likelihood phylogeny constructed from the genetic relationships of 173 BUSCO genes. One star on the module labels represents significant differentiation between populations.

Differences in metabolite expression between densities were evident for some metabolites. The aflatoxin backbone gene was most highly expressed in population A, particularly by NRRL3357 in low density (2mAO), although there was variation within the population. The expression of aflavarin, ATM1, AsparasoneA (all of which are core genes, Figure S9) and Piperazines (lnB) was high in populations B and C at low density, but not at higher density. Only isolates from population A showed medium-high expression of these genes at high density. The lovastatin-like gene cluster was significantly differentiated between populations at low density, but not at high density. Under both conditions, expression of this backbone was lower in *A. oryzae* and higher in population A.

### Population-specific SM BGCs

Because domesticated organisms often retain gene content inherited from their ancestral populations, we examined the SM BGCs of *A. oryzae* in the context of broader *A. flavus* population diversity. We focused on differences between *A. oryzae* and *A. flavus* populations as comparisons within *A. flavus* populations were previously characterized by Drott et al. (2021).

*A. oryzae* retains nearly all BGCs found across *A. flavus* populations, reflecting its origin from within a large, shared core pangenome. Consistent with previous work, we do not find pervasive large-scale deletions of the aflatoxin gene cluster in *A. oryzae* (although some are evident), suggesting that inability to produce aflatoxin is often the result of SNPs, smaller indels, or regulatory changes. A subset of BGCs is found in low frequency or absent in *A. oryzae*, but these clusters are generally not fixed in the broader *A. flavus* population (e.g., BGC_5, Figure S9).

Interestingly, some population-specific BGCs appear in *A. oryzae* even when absent from other lineages. For example, the astellolide cluster, also found in population C, is present in some *A. oryzae* isolates. In other cases, population-specific BGCs occur at higher frequency in *A. oryzae* than in the population from which they were first observed (e.g., BGC_4 – see BGC_8, BGC_4, Figure S9).

Several novel BGCs appear to be unique to *A. oryzae* but all of these occur at very low frequencies. The cluster dubbed ‘combo99’, for example, which shares 62% identity with auerobasidin A1 backbone across 73% of the protein from *Aureobasidium pullulans*, is found in only a few *A. oryzae* genomes. Additionally, rare clusters that were previously excluded from population-level analysis of Drott et al. 2021 as singletons— such as combo100 and combo101—are now included due to their presence in multiple *A. oryzae isolates*.

### Gene regulatory networks reveal differentiated transcriptomic architectures

To understand regulatory influences on transcriptional differences across conditions and populations, we constructed gene regulatory networks using significantly differentially expressed genes. Networks constructed with DEGs from condition and condition * population contrasts were small (Figure S11). In the density network, only one regulator, 5739 (Aspfl2_3 genome), showed high centrality, targeting genes from BGC_90 and BGC_51 (Figure S11B). In the condition * population contrast, 2210367 regulated 2207763 (BGC_47) and transcription factor 2283259 (BGC_52).

The population-based network included 1,978 nodes. The highest-centrality regulators were not strongly associated with BGCs (see central-right mass in Figure 4). For example, 2225371, the top-ranked transcription factor, regulated 158 genes, but only two (from BGC_67 and BGC_89) were BGC-associated. Regulators 635, 8311, and 2069970 shared substantial target overlap, together regulating 393 genes, only three of which were in BGCs. This suggests large-scale population transcriptomic shifts are driven by regulators of primary rather than secondary metabolism. Supporting this, the global secondary metabolism regulator LaeA (1256857) was not differentially expressed.

**Figure 4.**
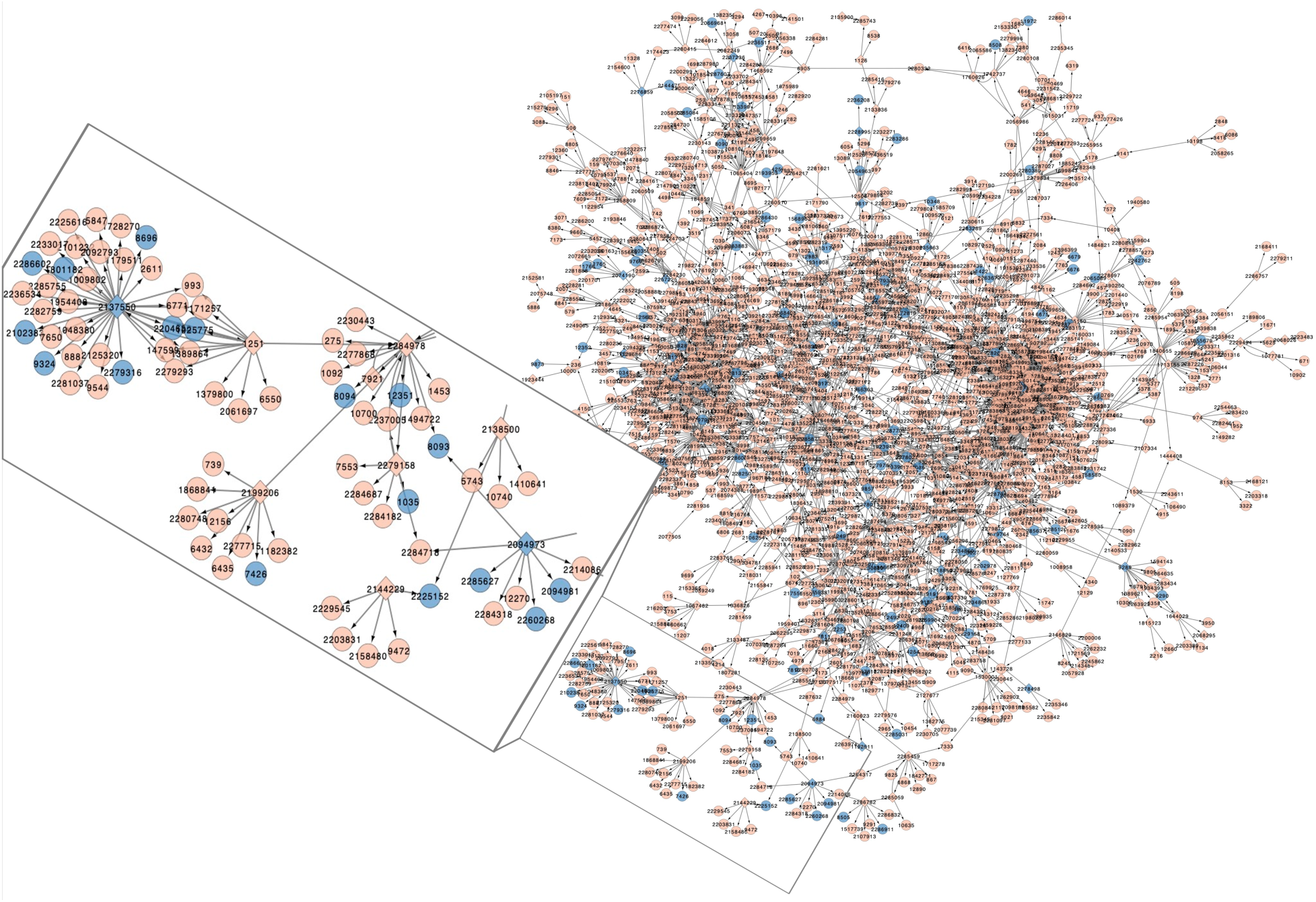
Gene regulatory network of genes that are significantly differentially expressed between *Aspergillus flavus* populations (A, B, C, and *Aspergillus oryzae*). Nodes represent genes and are labeled with protein IDs from the NRRL3357 JGI Aspfl2_3 genome. Nodes corresponding to transcription factors are presented as diamonds while circles represent all other genes. Genes located in a secondary metabolism gene cluster predicted in the reference genome are colored blue. The size of nodes indicates the IVI centrality index, with larger nodes having higher centrality scores. Arrows indicate inferred regulatory direction. A portion of the network that shows regulatory interactions between BGCs is enlarged for clarity and is discussed in the text. A small number of genes from this network that shared no linkage with the main network displayed here were omitted for visual clarity but can be found in Figure S10.

In contrast, several lower-centrality transcription factors showed targeted regulation of secondary metabolism. Notably, 2137550 (AFL2G_00934.2, from the asparasone A cluster) appears to regulate its own cluster backbone gene (2279316) in cis and two aflatrem cluster genes—including the backbone (2204636)—in trans. It also targets genes from the piperazine cluster (1801182) and BGCs 35, 44, 57, and 85 (non-backbone; Figure 4). The aflatrem backbone is also co-regulated by 1251 (AFLA_084720), and both 2137550 and 1251 coregulate a hypothetical gene in BGC85, with the former also regulating 1251, suggesting potential regulatory feedback.

Additional cis- and trans-BGC regulation included transcription factor 2094973 from the lovastatin-like cluster, which regulated three cluster genes (a P450, MFS transporter, and FAD monooxygenase; see callout in Figure 4). This TF also putatively co-regulates a reductase in BGC24 with 2284978 (AFLA_054800), which is itself downstream of 1251 and 2282289. Interestingly, 2282289 may also regulate the TF in the ustiloxin B cluster, 1718850, which controls six of the cluster’s genes in a population-specific way where *A. oryzae* is marked by lower expression. These results suggest cluster-specific transcription factors sometimes mediate trans-BGC expression, as has been found in several fungal species (Bergmann et al., 2010; Shostak et al., 2020).

## Discussion

The relationship of modern domesticated organisms to ancestral populations of wild relatives has clarified the evolution of dogs (Bergström et al., 2022), silkworms (Tong et al., 2022), and red beans (Chien et al., 2025), helping to elucidate which traits evolved through domestication. Similarly, our results suggest that hallmark features of *A. oryzae* may have begun evolving prior to domestication. Aflatoxin production is reduced in populations C (Drott et al., 2020), and the genes in this cluster are sometime degraded or altogether lost in isolates of population D (Hatmaker et al., 2025); both of these populations share more recent common ancestry with *A. oryzae*, raising the possibility that some softening of mycotoxigenic traits preceded domestication. While the geographic distribution of population D remains unclear, population C is enriched in northern latitudes (Drott et al., 2020) that align with the historical context of *A. oryzae* domestication. Transcriptomic signatures of domestication are also evident: *A. oryzae* shows the strongest expression of module 14, enriched in carbohydrate metabolism, with population C following closely. Notably, *A. oryzae* retains the capacity to express many secondary metabolite genes, sometimes mirroring expression patterns seen in other populations.

*Aspergillus oryzae* has long been considered safe and non-pathogenic due to the lack of aflatoxin production (Barbesgaard et al., 1992). We were therefore surprised to find that *A. oryzae* isolates fall within a clade of *A. flavus*—population D—enriched in clinical strains (Hatmaker et al., 2025). *A. oryzae* retains the ability to infect animals, though with lower virulence than *A. flavus* in a zebrafish model, consistent with past studies (Barbesgaard et al., 1992; Ford & Friedman, 1967). Although putative *A. oryzae*-infected samples have been identified from patients (Barbesgaard et al., 1992), these isolates may have been misidentified. Difficulties differentiating *A. oryzae* from *A. flavus* have been lamented in the literature for over a century (Thom & Church, 1921; Wehmer, 1901). Our results clarify some of the confusion: while *A. oryzae* has previously been reported as monophyletic (Watarai et al., 2019), suggesting a single domestication event, the inclusion of population D isolates resolves *A. oryzae* as polyphyletic. The inclusion of the *A. flavus* type strain, NRRL1957, in the polyphyletic distribution that includes many industrial isolates argues for a true polyphyly. While A. oryzae could escape from industrial facilities, intermingling with environmental strains, the high clonality of industrial isolates and relatively diversity of clinical/environmental isolates does not support this as an explanation for the patterns we observe. Multiple domestications (and/or post-domestication admixture) may better align with historical accounts of ‘spontaneous’ fermentation practices (Allwood et al., 2023). More work is needed to define the boundaries of *A. oryzae* within population D, and although it may share ancestry with the clinically enriched population D, our findings do not indicate increased risk.

Domestication has likely reshaped the ecology of *A. oryzae* by altering its nutrient landscape and promoting growth at unnaturally high densities. These shifts may help explain distinct gene expression patterns observed in domesticated strains -- for instance, the overrepresentation of module 14, enriched for carbohydrate metabolism -- mirroring findings from earlier studies (Gibbons et al., 2012) and a trait involved in traditional A. oryzae food fermentations (Han et al., 2024). While the expression of many SMs varied between populations (Figure 3), *A. oryzae* retained the ability to express many SM genes, emphasizing that these genes are not entirely pseudogenized. Although transcriptomic profiles do not always directly correlate with metabolite production (Bergmann et al., 2010; Shostak et al., 2020), many of our findings align with previously reported phenotypes, and transcriptomes provide insight into which pathways are actively engaged. Many transcriptomic differences identified here extend beyond those found in domesticated *A. oryzae*, suggesting that large-scale reorganization of gene expression is a general mechanism of microbial adaptation. While fungal evolutionary studies have often avoided focus on transcriptomic divergence -- perhaps due to concerns about its heritability -- bacterial systems increasingly recognize these traits as both adaptive and sometimes genetically encoded (Ryall et al., 2012). The population-level differences were often consistent across individuals, supporting a model in which heritable regulatory variation contributes to ecological specialization. One striking example is module 22, associated with development and reproduction, which is markedly downregulated in population B—a group with limited recombination and a skewed conidia-to-sclerotium ratio (Drott et al., 2020). These findings position population-specific transcriptomic architectures as a powerful, and underappreciated, lens for studying fungal ecology and evolution.

Population transcriptomic differentiation in *Aspergillus* may confer adaptive flexibility in response to ecological shifts. We examined the effects of colony density— an environmental parameter central to industrial *koji* production and relevant in natural contexts where spore dispersal varies across spatial scales. Prior work has linked density to shifts in aflatoxin and sclerotial production via oxylipin signaling (Affeldt et al., 2012). Our results extend this framework by revealing substantial, population-specific transcriptomic rewiring in response to density. For example, while NRRL3357 (2mAO) exhibited strong density-dependent induction of aflatoxin genes, this pattern was not consistent across other isolates. Notably, three SM clusters—asperasone A, aflatrem, and aflavarin, all associated with sclerotia—were strongly induced at low densities in populations B and C, but not in *A. oryzae*. While population B is known for prolific sclerotial development, population C typically forms few or no sclerotia (Drott et al., 2020) suggesting that activation of these clusters may not strictly depend on structure formation. This potential decoupling raises questions about other structure-related compounds like xanthones in *A. nidulans* cleistothecia (Liu et al., 2021). Our findings underscore the importance of considering transcriptional plasticity not only as a downstream consequence of development, but as a dynamic population-specific ecological strategy.

The genetic regulation of secondary metabolism in *Aspergillus* has been widely studied, revealing that BGCs often share co-localized cis-BGC regulatory elements conserved across evolutionary timescales (Rösler et al., 2016; Seong et al., 2009; Woloshuk et al., 1994). Global regulators such as *laeA* orchestrate broad transcriptional control of multiple SMs within a genome (Bok & Keller, 2004). However, most of this work has relied on single reference strains, offering limited insight into how regulatory architectures vary within species. Our population-level gene regulatory networks suggests that differences in both secondary and primary metabolism can arise from differentially regulated transcription factors (TFs), rather than changes in core BGC architecture. Notably, we identify evidence for both cis-BGC-acting regulation (e.g., *ustiloxin*) and rare trans-BGC regulation, as seen in a piperazine-associated TF that appears to control multiple unrelated clusters. Although trans-regulatory influences across BGCs are seldom reported, exaptation of regulators has been observed (Bergmann et al., 2010; Shostak et al., 2020; Wang et al., 2021), emphasizing the potential for dynamic interactions between BGC regulators. While further validation is needed to confirm the relationship of specific transcription factors to population-specific transcriptomic patterns, these findings suggest that transcriptomic restructuring between populations occurs at a modular scale impacting the expression of one or several metabolites at a time while *laeA* remains stable.

In *A. flavus*, gene regulation, particularly of secondary metabolism, has population-specific responses to ecological context, including colony density. Canonical features of *A. oryzae* domestication like increased carbohydrate metabolism and low aflatoxin production may have begun evolving before domestication; after domestication, *A. oryzae* retains expression of some SMs, at least at the transcript level, and virulence. The distribution of *A. oryzae* isolates suggests multiple domestication events, or admixture. By leveraging population-scale gene expression and inferred regulatory networks, we uncover modular rewiring of population-specific SM pathways, suggesting that cis and trans regulation have both contributed to diversification. Population-specific responses to environmental stimuli could have significant implications for fungal evolution and our understanding of mycotoxigenic threats in agriculture. These results offer a framework for understanding how domestication and population ecology interact to shape fungal phenotypes and raise questions about the evolutionary histories of industrial fungi like *A. oryzae*.

## Acknowledgements

E.A.H. was supported by a postdoctoral fellowship funded by the USDA Agricultural Research Service’s SCINet Program and AI Center of Excellence (ARS project numbers 0201-88888-003-000D and 0201-88888-002-000D) and administered by the Oak Ridge Institute for Science and Education (ORISE) through an interagency agreement between the U.S. Department of Energy (DOE) and the U.S. Department of Agriculture (USDA). ORISE is managed by the Oak Ridge Associated Universities (ORAU) under DOE contract number DE-SC0014664. M.T.D. and JME received support from USDA-ARS project 5062-21220-024-000D. Research in A.R.’s lab is supported by the National Science Foundation (DEB-2110404) and the National Institutes of Health/National Institute of Allergy and Infectious Diseases (AI153356). NMMS was supported on the NSF-GRFP award DGE-2137424. All opinions expressed in this paper are the authors’ and do not necessarily reflect the policies and views of USDA, DOE, or ORAU/ORISE. J.G.G.’s contribution was funded through NSF Grant 1942681.

## Data availability

All transcriptomic and genomic data generated here will be made available through NCBI upon acceptance of this manuscript for publication.

## Conflicts of interest

AR is a scientific consultant for LifeMine Therapeutics, Inc. All other authors declare no competing interests.

