## Supplementary Methods and Figures for "Population-Specific Transcriptomic Shifts Underlie Secondary Metabolic Diversification in *Aspergillus flavus* and the Domestication of *Aspergillus oryzae*"

### Supplemental Methods:

BUSCO Maximum likelihood phylogeny construction:

Genes were aligned using the ' L-INSI-i' model for 177 gene dataset in MAFFT (Katoh et al., 2002) parameter 'automated1' parameter in trimAL (Capella-Gutiérrez et al., 2009). Attempts to use higher-accuracy alignment approaches (e.g., L-INSI-i in MAFFT), were computationally prohibitive across the larger BUSCO dataset. To avoid potentially spurious alignments, we removed all genes where alignment lengths differed by 50% or more compared to the unaligned and untrimmed lengths similar to previous work (Steenwyk et al., 2019). Resulting alignments were used to construct into a maximum likelihood phylogeny using an automated partition model selection and 1,000 ultra-fast bootstraps in IQ-TREE2 (Nguyen et al., 2015)

BUSCO quartet-based phylogeny construction:

Phylogenetic trees were also constructed for each of the genes in the Eurotiales BUSCO dataset using the 'auto' parameter in MAFFT (Katoh et al., 2002) parameter 'automated1' parameter in trimAL (Capella-Gutiérrez et al., 2009). These trees were used to construct a quartet-based tree in ASTRAL (Zhang et al., 2018). Because ASTRAL's computational load is orders-of magnitude less than maximum-likelihood trees, the quartet-based analysis was run on the Euratiales BUSCO set.

SNP maximum likelihood phylogeny construction:

SNPs were subset to retain only biallelic sites that passed filters for the full dataset. This dataset was also subset to remove all SNPs that fell within genes (as these are more likely to be subject to selection) and to retain only SNPs that were at least 1,000 bp apart (to increase independence) using vcftools v0.1.16 (Auton and Marcketta 2009). These datasets were exported to nexus and fasta formats using the script vcf2phy (Ortiz, 2019) and constructed into neighbor-net networks using SplitsTree (Huson, 1998) and maximum-likelihood trees using IQ-TREE2 (Nguyen et al., 2015) with parameters identical to those defined above.

SNP maximum likelihood phylogeny construction with Hatmaker Data:

We also incorporated *A. oryzae* data into a recent population-genomic analysis of *A. flavus* that identified a new population of *A. flavus* that emerged after this study had been designed (Hatmaker et al., 2025). In brief, SNPs called from mapping to NRRL3357 (GCF_009017415.1) from both *A. flavus* (n = 255) and *A. oryzae* (n = 58) were combined, genotyped, and filtered using GATK v4.1.8 and Picard tools v3.1.0. The resulting GVCF file of SNPs from 314 isolates (Table S3), including the outgroup *Aspergillus minisclerotigenes* (SRR12001146), was formatted as a PHYLIP file using the tool vcf2phylip.py (Ortiz, 2019). The script ascbias.py (https://github.com/btmartin721/raxml_ascbias) was used to remove invariant sites from the data for ascertainment bias correction. The phylogeny was built using IQ-TREE2 v2.2.2.6 (Stamatakis, 2014) under the "TVM+F" model with 1,000 replicates for ultrafast bootstrapping. The tree was visualized and annotated in iTOL v6 (Letunic & Bork, 2024).

**Supplemental Results:**

Pangenomic patterns

We constructed an *A. flavus* pangenome to understand the overlap in gene content between *A. flavus* populations and to facilitate our interpretation of the expression of genes of interest through copy-number variation of orthogroups. Extensive interpretation of *A. flavus* pangenomics has recently been completed by Hatmaker et al. (https://www.nature.com/articles/s41467-025-62777-9), and, given the focus of this study on transcriptomics, was beyond the scope of this manuscript. Our pangenome consisted of 15,913 orthogroups that were present in at least two individuals. Another 5,699 orthogroups were found in only a single individual. The openness of the pangenome depended on the inclusion of x and y groups, emphasizing that the predictive power of rarefaction curves is dependent on what populations are being sampled. Overall, the size of the pangenome and its openness within *A. flavus* populations is consistent with Hatmaker et al. (Hatmaker et al., 2025).

The length of genes decreases across the core, accessory, shell and cloud genome continuum. (figure 1 B). While the focus of this manuscript is secondary metabolism and transcriptomic rearrangements, we did perform some preliminary analyses of pangenomic enrichment. Only 63 interPRO terms were significantly differentiated between populations. Surprisingly, we did not find any enrichment of IPR006047, the term associated with alpha-amylase activity ( P = 0.752 [f isher_out.summary]). While past work has document duplications of certain alpha-amylase genes, our results suggest that on a population-scale the total number of genes with this activity has not changed, emphasizing that not all genes with this putative function necessarily have the same ecological functionality. A series of terms associated with viral reproduction (IPR000477, IPR041373, and IPR001584) were significantly enriched among Oryzae isolates, raising questions about the ability of domesticated isolates to purge viral sequences. Interestingly an enrichment of retrotransposon gag proteins (IPR005162) was also found in population B, a population previously shown to have relatively limited evidence of recombination. Terms associated with chitin recognition (IPR001002), starship-transposable elements (IPR021842), and zinc binding (IPR041588) were also significantly enriched in Oryzae. The only significantly enriched term that was overtly related to secondary metabolism was IPR019587 (polyketide cyclase/dehydrase activity) which was found in highest abundance in population A.

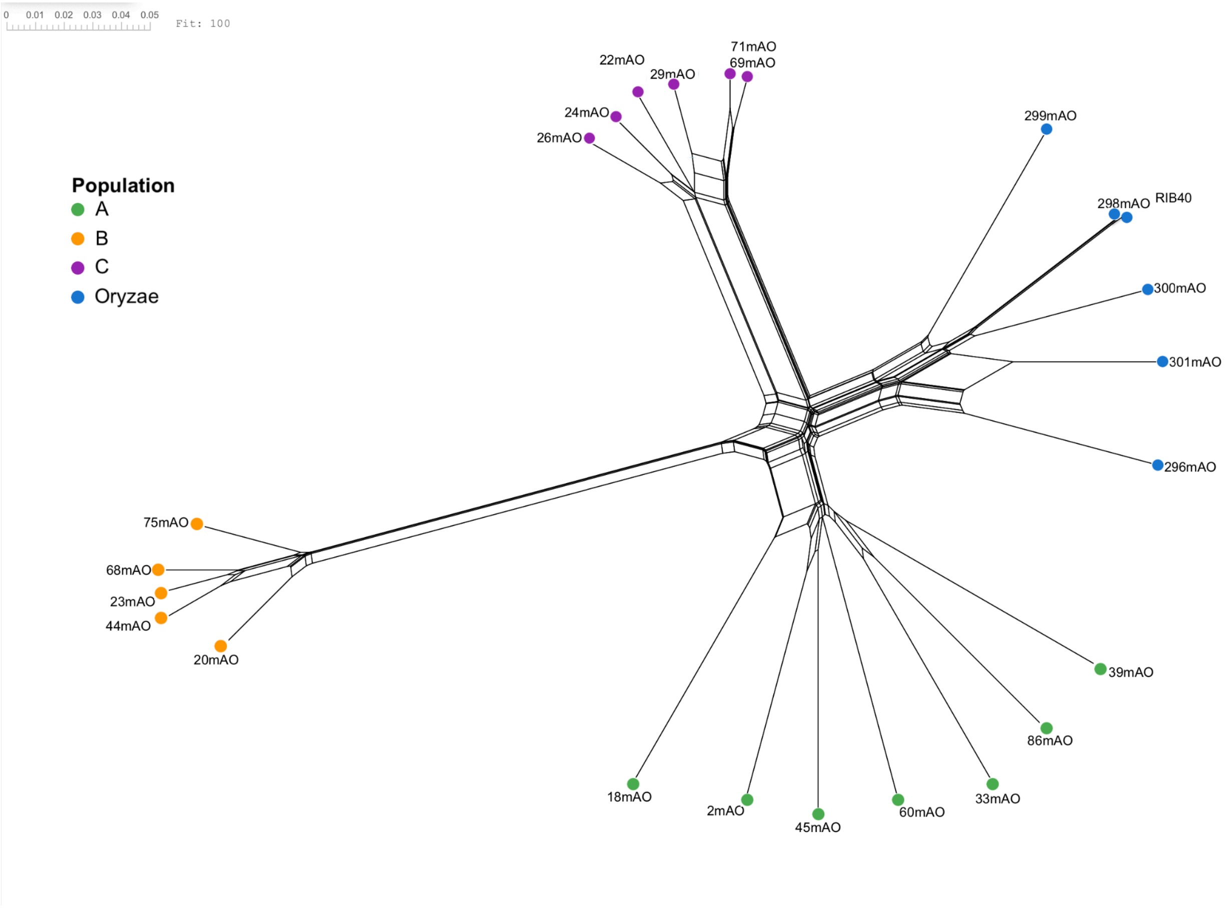


Figure S1. Neighbor-net network constructed in SplitsTree from 793,820 biallelic SNPs.

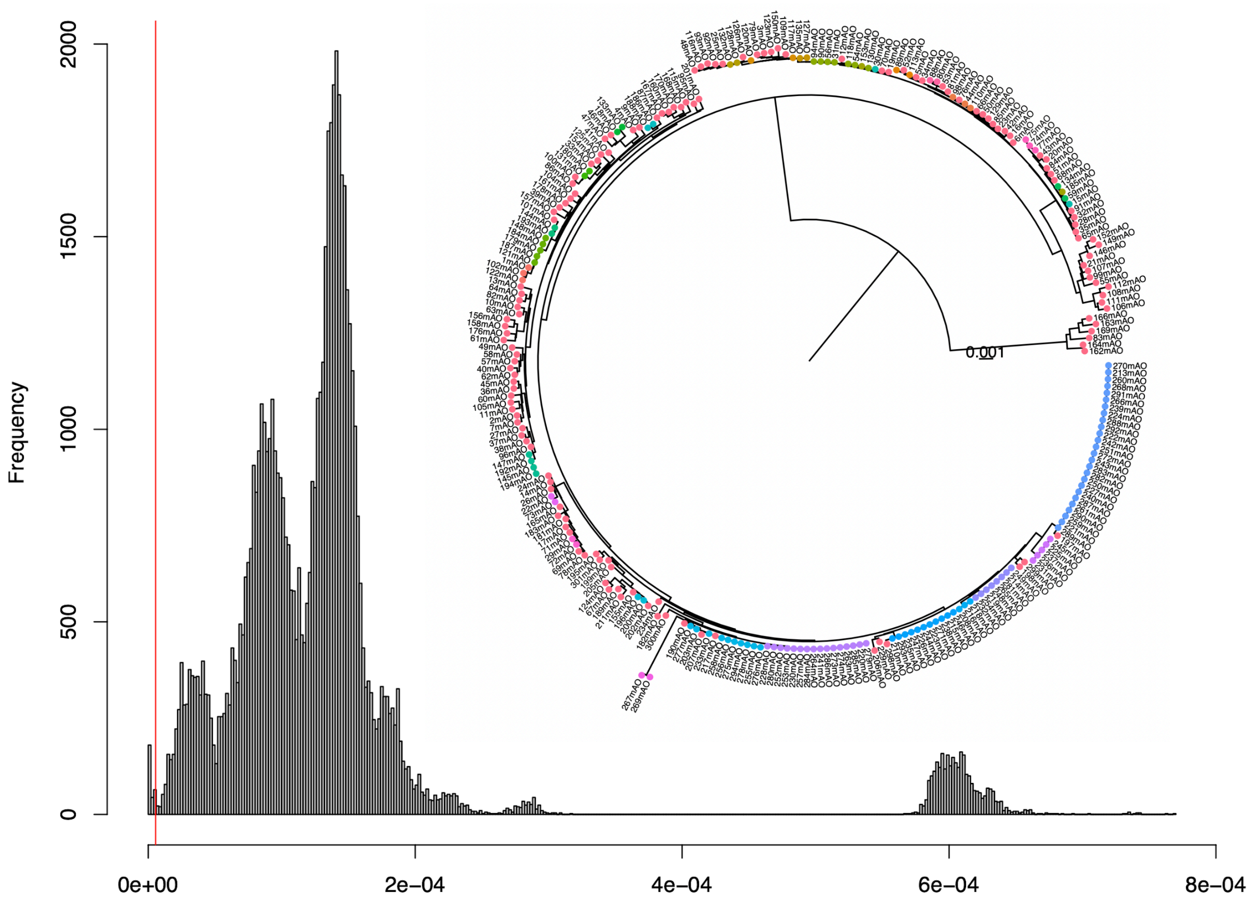


Figure S2. Clone correction of *Aspergillus flavus* populations was achieved by selecting a genetic similarity cutoff (indicated with a vertical red line) that separated a distribution of very closely related isolates as inferred using a histogram of genetic distances inferred from a maximum likelihood phylogeny of 177 BUSCO genes (Figure S1). The resulting assignments of multi-locus genotypes (MLGs) was visualized onto Figure S1 where MLGs with only one representative are all colored the same shade of red, but all MLGs with more than one representative are colored to match only their corresponding clones.

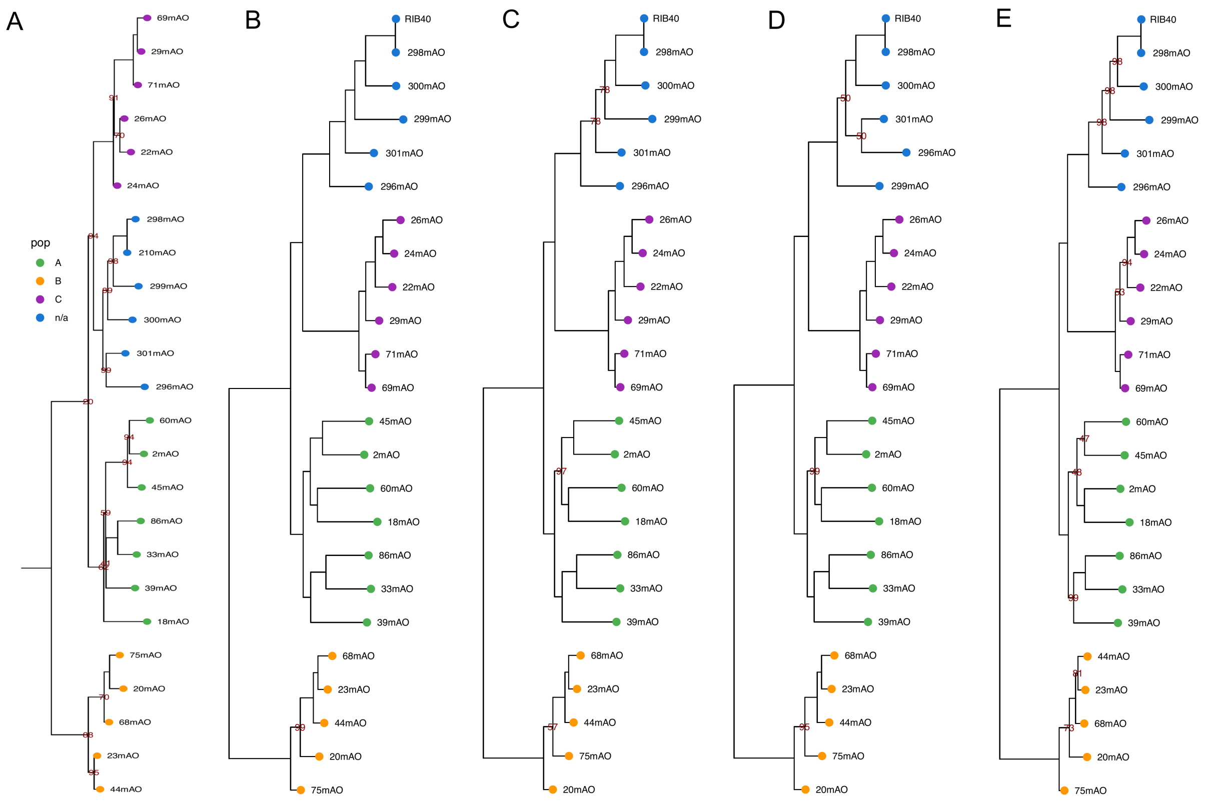


Figure S3. Maxmimum likelihood phylogenies of a subset of isolates used for wet-lab experiments in this study. Trees were assembled from 177 high quality BUSCO genes (A), all 793,820 biallelic SNPs (B), 629,271 biallelic SNPs with no missing data (C), 430,178 SNPs that were outside of genes (minimizing the impact of selection) (D), and 20,067 SNPs found outside of genes and thinned to a least 1kb (minimizing selection and maximizing marker independence) (E). Bootstraps calculated from 1,000 ultrafast bootstraps in iqtree2 are indicated at nodes when they are less than 100.

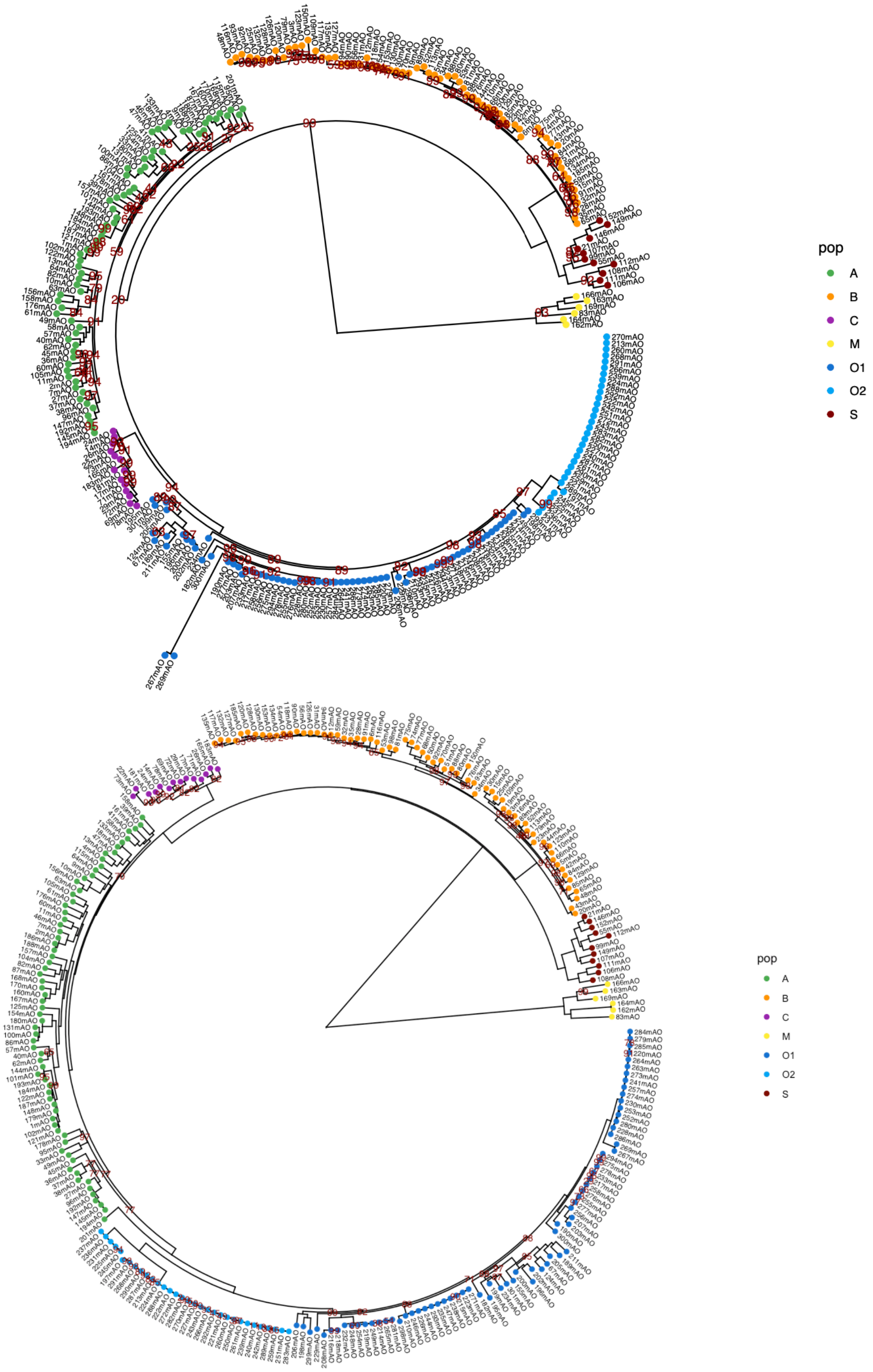


Figure S4. Maximum-likelihood phylogeny inferred from the nucleotide sequences of 177 Fungal (top) or 2,205 Eurotiales (bottom) high-quality BUSCO genes. Bootstraps calculated from 1,000 ultrafast bootstraps in iqtree2 are indicated at nodes when they are less than 100.

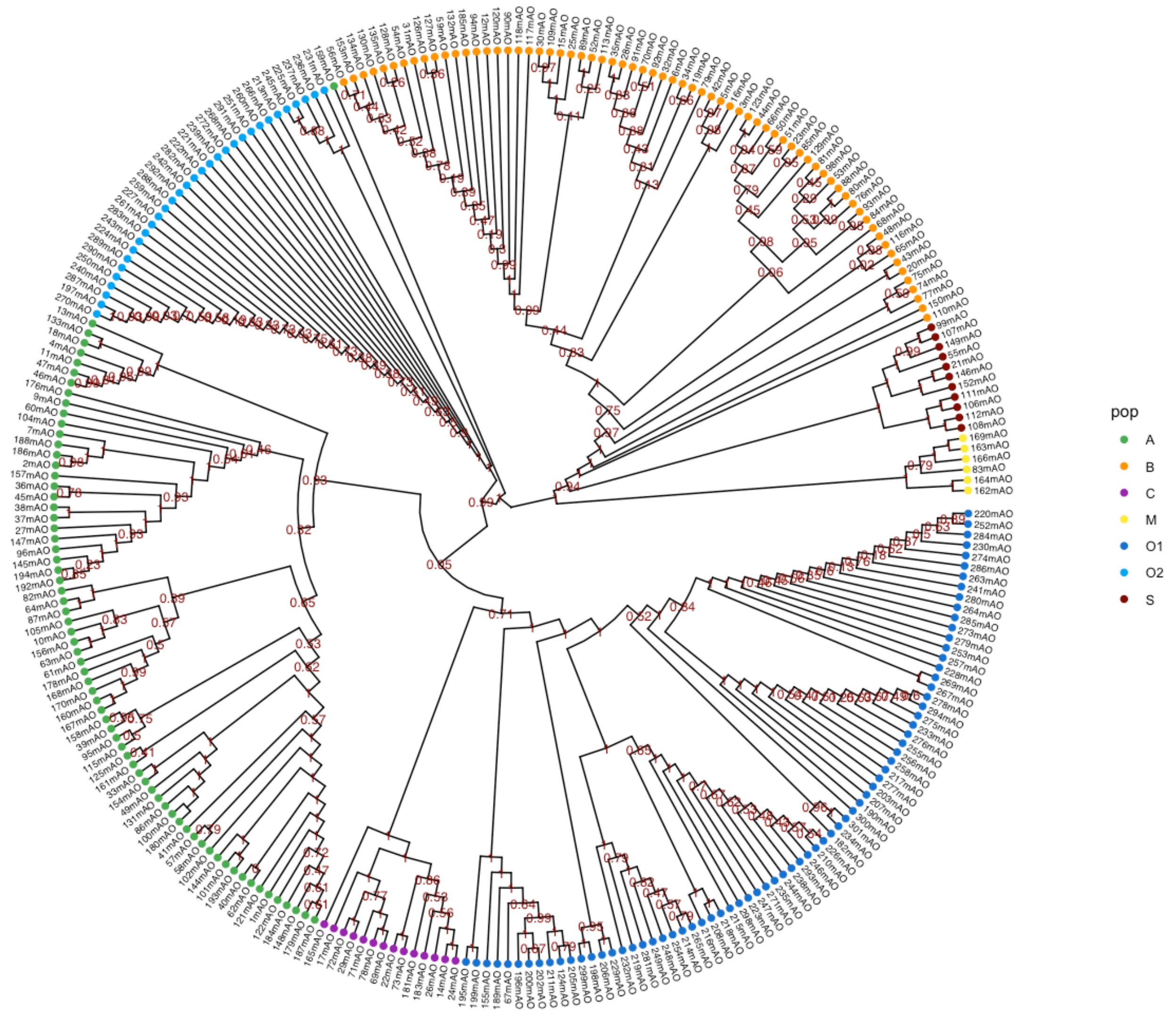


Figure S5. Astral cladogram constructed from 2,205 phylogenies made with nucleotide alignments of high-quality Eurotiales BUSCO genes. Posterior probabilities are indicated at each node. Tree is presented as a cladogram to allow for readability of posterior probabilities.

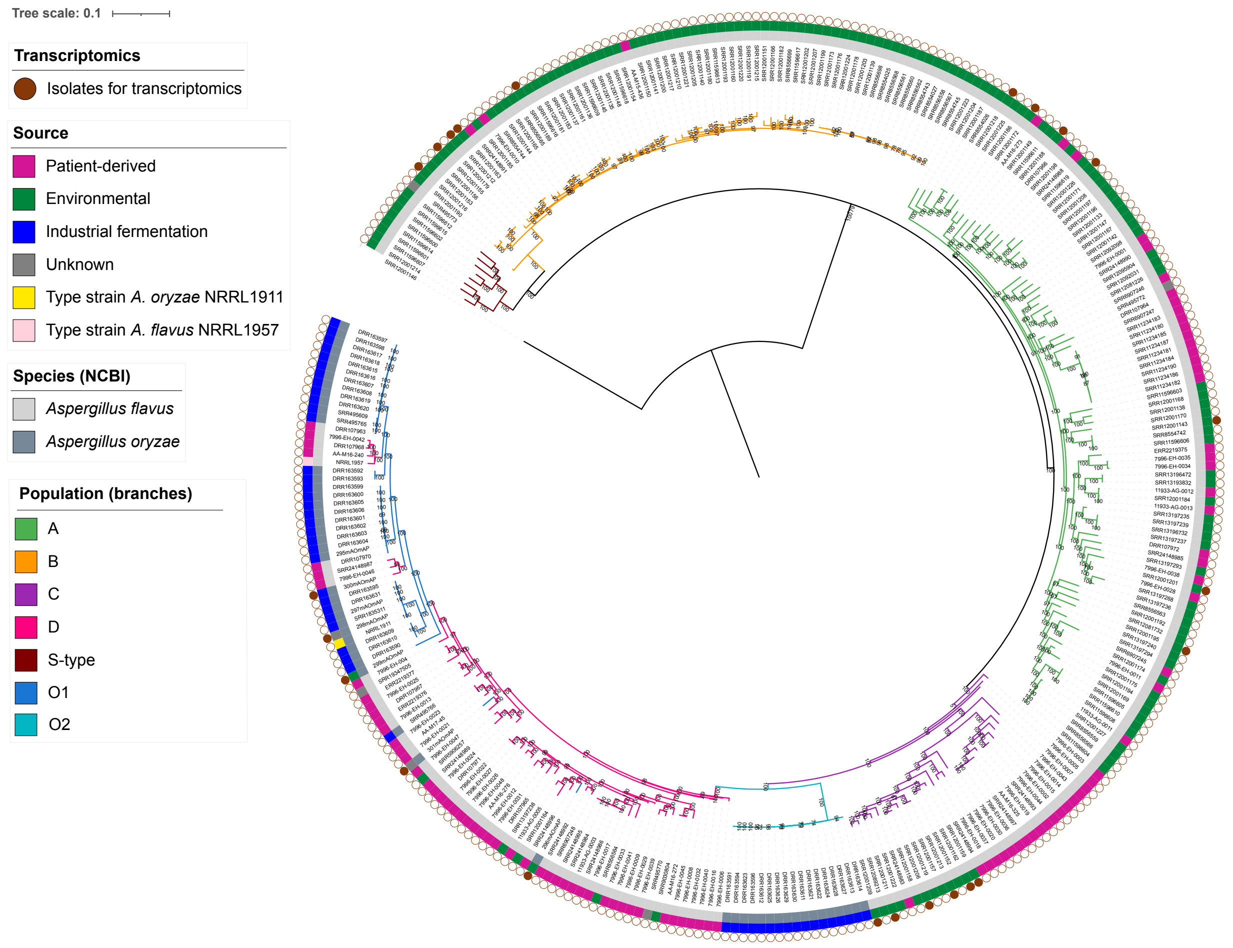


**Figure S6:** A maximum likelihood phylogeny constructed using genome-wide SNPs including data from Hatmaker et al. (Hatmaker et al., 2025) is rooted at *Aspergillus minisclerotigenes*. Bootstrap support values are the result of 1,000 ultra fast bootstrapping iterations.

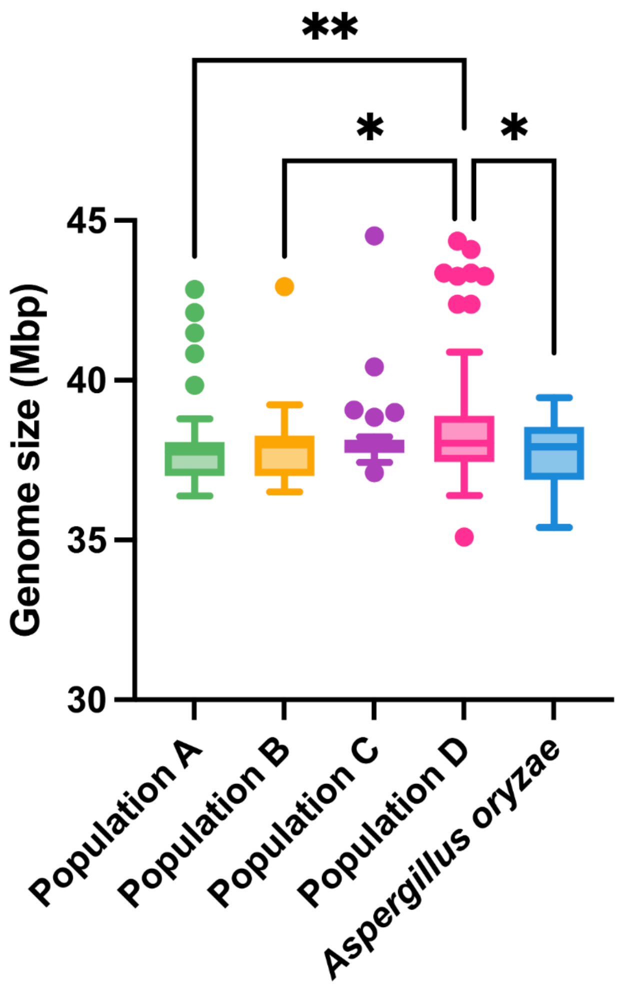


**Figure S7.** Genome size for *Aspergillus flavus* population D is significantly higher than *Aspergillus oryzae* isolates. Box and whisker plots show the genome size by population. The whiskers include the highest and lowest values within 1.5x the interquartile range (IQR), with individual dots representing isolates outside the 25th percentile minus 1.5x the IQR or 75th percentile plus 1.5x the IQR. *Aspergillus flavus* populations are color coded as follows: Population A in green (n = 82), population B in yellow (n = 69), population C in purple (n = 29), population D in pink (n = 72). *Aspergillus oryzae* is indicated in blue (n = 28). Asterisks indicate significance (one-way ANOVA with multiple comparisons; *p < 0.05, **p < 0.005), with all other comparisons not significant.

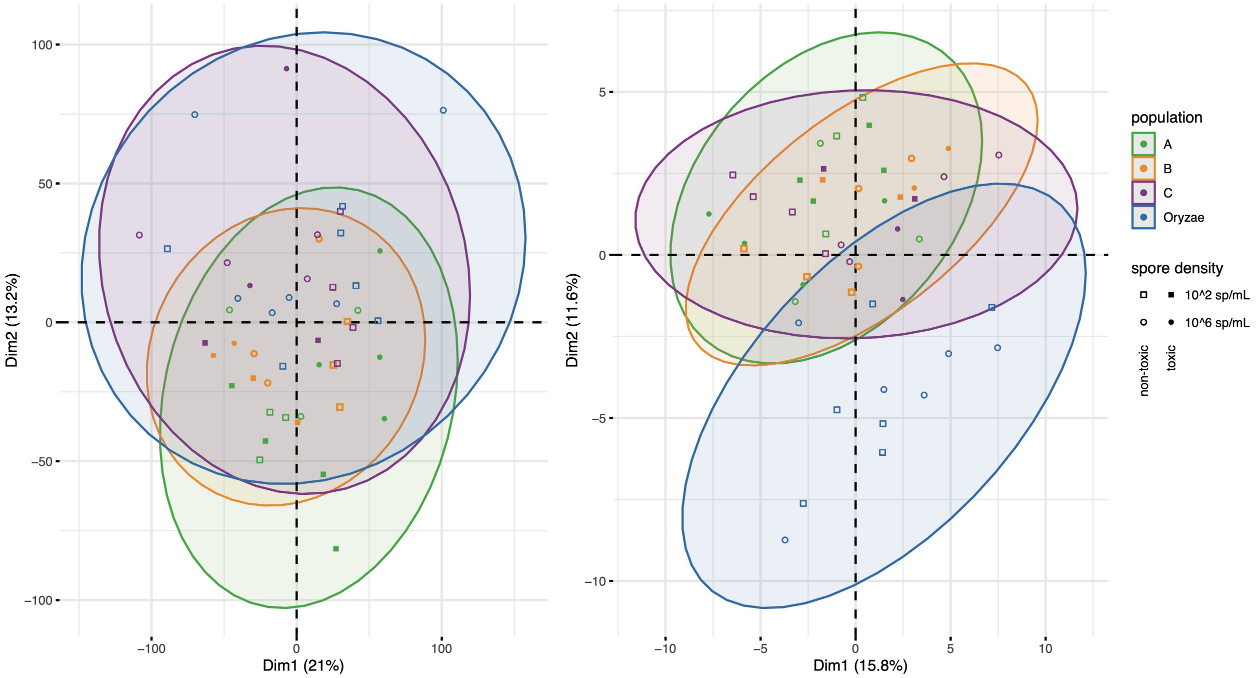


Figure S8. Principal component analysis of gene expression data for all genes (left) and secondary metabolite backbone genes (right) as a function of *Aspergillus flavus* population and spore density.

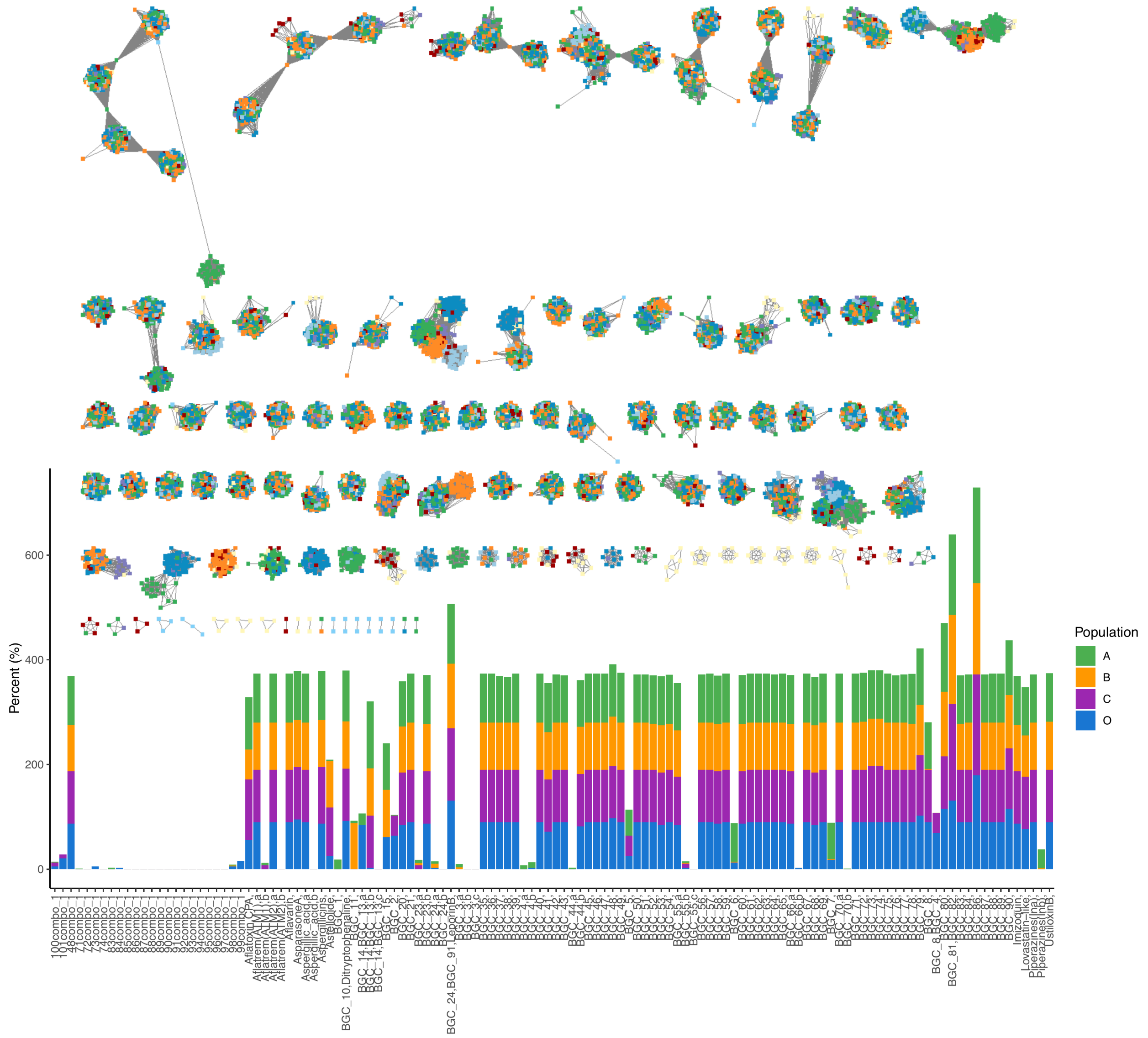


Figure S9. Analysis of secondary metabolite gene clusters found by antiSMASH in all genomes used in this study. Clusters have been networked into gene cluster families representing clusters likely encoding identical or similar compounds using BiG-SCAPE (top) where nodes represent a single gene cluster in a single genome. All colors represent population-level designations with dark red being Aspergillus flavus S-type and yellow being *Aspergillus minisclerotigenes*. The network represents all data including clones and includes a lighter blue color designating the A. oryzae clade O2 (see Figure S1). The network was dereplicated to separate out GCFs connected by few connections (i.e., bridge clusters) and clone corrected to calculate the percent of genetic individuals from each population that have a single gene cluster (stacked bar chart bottom). The bar chart in the bottom does not include S-type or A. minisclerotigenese; these groups entirely explain BGCs with empty bars. GCFs that overlap in physical space of the reference genome with results from Drott et al. (2021) have been renamed accordingly. However, we did not perform the extensive manual vetting of Drott et al. (2021), and in rare instances, some GCFs identified here overlap with several distinct clusters from Drott et al. (2021). In instances where our GCFs overlapped with more than one cluster from Drott, all overlapping clusters are listed in a comma separated list. Conversely, our dereplication pipeline occasionally separated a single GCF from Drott et al. (2021) into multiple GCFs, these have been designated with “a,b,c etc.” – this type of error often reflected slightly divergent versions of the cluster found in S-type and A. minisclerotigenese. A set of GCFs that did not map to results of Drott et al. are labeled as “combo”. While most of the ‘combo’ GCFs were found in only in A. oryzae and/or a single isolate of another A. flavus population (and thus not included in Drott et al. 2021), 48combo is found in most isolates across populations including the NRRL3357 reference and corresponds to a beta lactam cluster with the backbone gene “AFLA_084630” – this appears to have been overlooked in Drott et al. 2021 but does not change any conclusions therein. Population designations are inferred from phylogenies in ways that are appropriate for large-scale inferences of population specific patterns but should not be overinterpreted when only a single or small number of individuals are represented in a GCF.

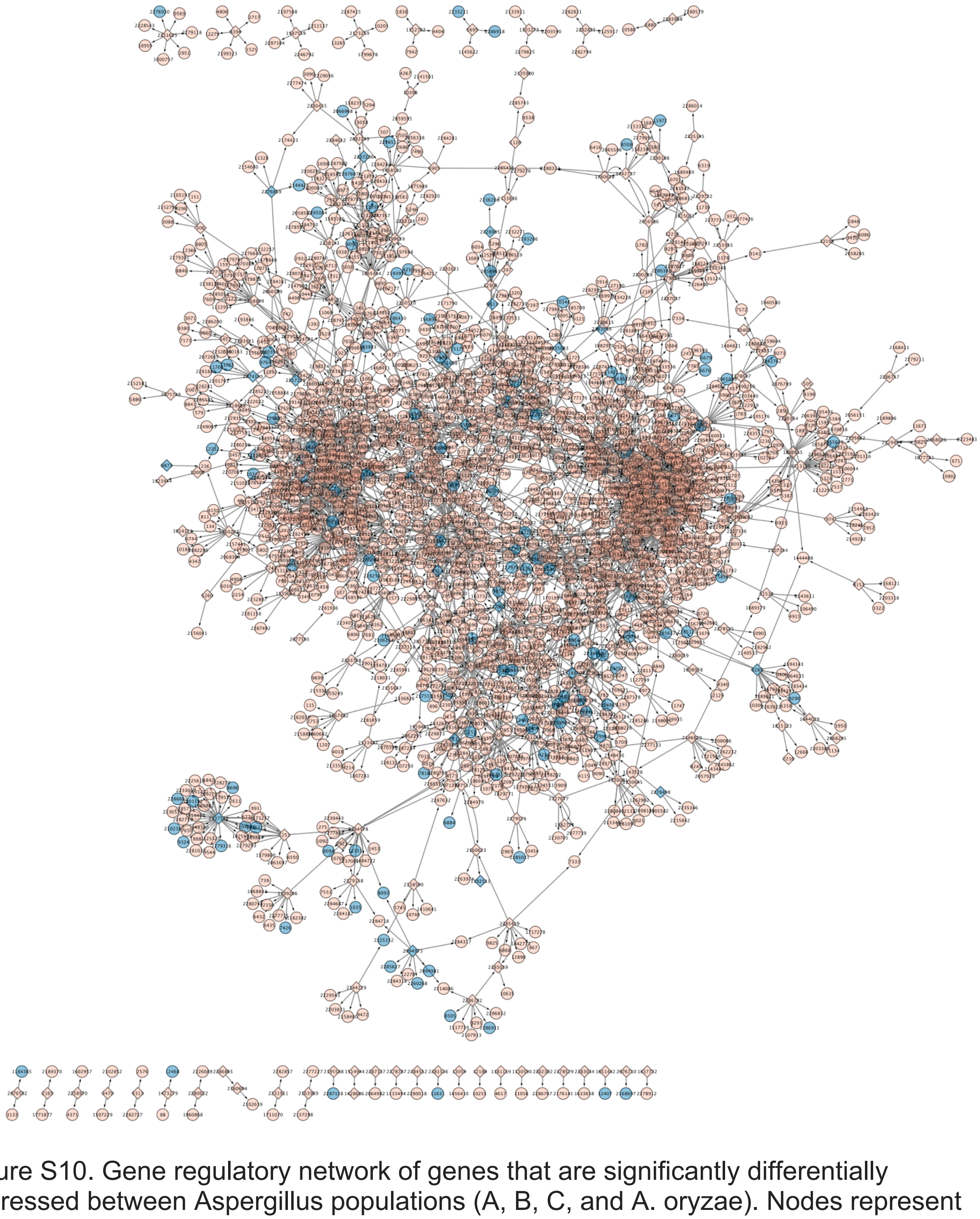

Figure S10. Gene regulatory network of genes that are significantly differentially expressed between Aspergillus populations (A, B, C, and A. oryzae). Nodes represent genes and are labeled with protein IDs from the NRRL3357 JGI reference genome. Nodes corresponding to transcription factors are presented as diamonds while circles represent all other genes. Genes located in a secondary metabolism gene cluster predicted in the reference genome are colored blue. The size of nodes indicates the IVI centrality index, with larger nodes having higher centrality scores. Arrows indicate inferred regulatory direction. A portion of the network that shows regulatory interactions between BGCs is enlarged for clarity and is discussed in the text. Node labels correspond to protein IDs from the JGI reference genome.

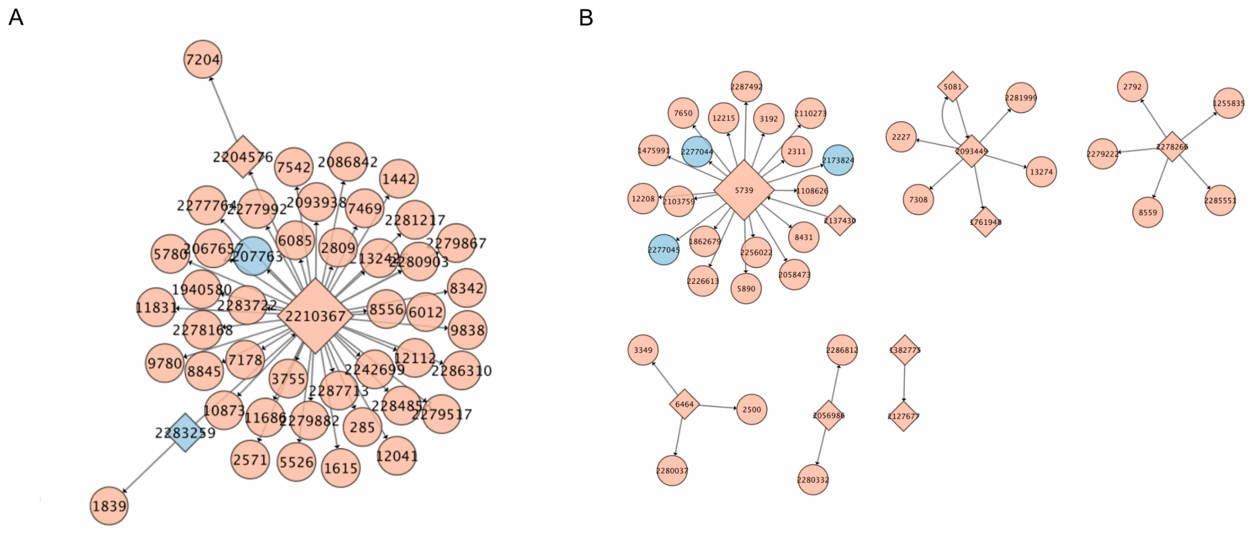


Figure S11. Gene regulatory network of genes that are significantly differentially expressed between Aspergillus populations (A, B, C, and A. oryzae) as a function of density (A) and those that differentiated solely between densities (B). Nodes represent genes and are labeled with protein IDs from the NRRL3357 JGI reference genome. Nodes corresponding to transcription factors are presented as diamonds while circles represent all other genes. Genes located in a secondary metabolism gene cluster predicted in the reference genome are colored blue. The size of —nodes indicates the IVI centrality index, with larger nodes having higher centrality scores. Arrows indicate inferred regulatory direction. A portion of the network that shows regulatory interactions between BGCs is enlarged for clarity and is discussed in the text. Node labels correspond to protein IDs from the JGI reference genome.

Capella-Gutiérrez, S., Silla-Martínez, J. M., & Gabaldón, T. (2009). trimAl: A tool for automated alignment trimming in large-scale phylogenetic analyses. *Bioinformatics*, *25*(15), 1972–1973.

Hatmaker, E. A., Barber, A. E., Drott, M. T., Sauters, T. J., Gumilang, A., Alastruey-Izquierdo, A., Garcia-Hermoso, D., Eagan, J. L., Keller, N. P., & Kontoyiannis, D. P. (2025). Population structure in a fungal human pathogen is potentially linked to pathogenicity. *Nature Communications*, *16*(1), 7594.

Huson, D. H. (1998). SplitsTree: Analyzing and visualizing evolutionary data. *Bioinformatics (Oxford, England)*, *14*(1), 68–73.

Katoh, K., Misawa, K., Kuma, K., & Miyata, T. (2002). MAFFT: a novel method for rapid multiple sequence alignment based on fast Fourier transform. *Nucleic Acids Research*, *30*(14), 3059–3066.

Letunic, I., & Bork, P. (2024). Interactive Tree of Life (iTOL) v6: Recent updates to the phylogenetic tree display and annotation tool. *Nucleic Acids Research*, *52*(W1), W78–W82.

Nguyen, L.-T., Schmidt, H. A., Von Haeseler, A., & Minh, B. Q. (2015). IQ-TREE: a fast and effective stochastic algorithm for estimating maximum-likelihood phylogenies. *Molecular Biology and Evolution*, *32*(1), 268–274.

Ortiz, E. (2019). *convert a VCF matrix into several matrix formats for phylogenetic analysis. 2.0*.

Stamatakis, A. (2014). RAxML version 8: A tool for phylogenetic analysis and post-analysis of large phylogenies. *Bioinformatics*, *30*(9), 1312–1313.

Steenwyk, J. L., Shen, X.-X., Lind, A. L., Goldman, G. H., & Rokas, A. (2019). A robust phylogenomic time tree for biotechnologically and medically important fungi in the genera Aspergillus and Penicillium. *MBio*, *10*(4), 10–1128.

Zhang, C., Rabiee, M., Sayyari, E., & Mirarab, S. (2018). ASTRAL-III: polynomial time species tree reconstruction from partially resolved gene trees. *BMC Bioinformatics*, *19*, 15–30.
